# ARHGEF17/TEM4 regulates the cell cycle through control of G1 progression

**DOI:** 10.1101/2024.04.14.589449

**Authors:** Diogjena Katerina Prifti, Eva Calvo, Annie Lauzier, Chantal Garand, Romain Devillers, Suparba Roy, Alexsandro Dos Santos, Laurence Descombes, François Bordeleau, Sabine Elowe

**Affiliations:** Centre de recherche du Centre Hospitalier Universitaire (CHU) de Québec-Université Laval, Axe de réproduction, santé de la mère et de l’enfant, Québec, QC G1V 4G2, Canada; Centre de recherche du Centre Hospitalier Universitaire (CHU) de Québec-Université Laval, Axe de Cancer, Québec, QC G1V 4G2, Canada; PROTEO-regroupement québécois de recherche sur la fonction, l’ingénierie et les applications des protéines, Québec, QC G1V 0A6, Canada; Département de biologie moléculaire, biochimie médicale et pathologie, Faculté de Médecine, Université Laval, Québec City, QC G1V 0A6, Canada; Département de Pédiatrie, Faculté de Médicine, Université Laval, Québec City, QC G1V 0A6, Canada; Centre de recherche sur le cancer de l’Université Laval, Québec, QC G1R 2J6, Canada; Centre de recherche en organogénèse expérimentale de l’Université Laval (LOEX), Québec, QC G1J 1Z4, Canada

## Abstract

The Ras homolog (Rho) small GTPases, via their role in regulating the actin cytoskeleton, coordinate diverse cellular functions including cell morphology, adhesion and motility, as well as cell cycle progression, survival and apoptosis. The upstream Rho regulators for many of these functions are unknown. ARHGEF17 (also known as TEM4) is a Rho family guanine nucleotide exchange factor (GEF) that been implicated in cell migration, cell-cell junction formation and the mitotic checkpoint. In this study we characterize the regulation of the cell cycle by TEM4. We demonstrate that TEM4 depleted cells exhibit multiple defects in mitotic entry and duration, spindle morphology, and spindle orientation. In addition, we find that TEM4 insufficiency leads to excessive cortical actin polymerization and cell rounding defects. Mechanistically, we demonstrate that TEM4 depleted cells delay in G1 as a consequence of elevated levels of the G1/S inhibitor p21^waf1/cip1^ and that TEM4 depleted cells that progress through to mitosis, do so with decreased transcription of *CCNB1* and thus attenuated levels of cyclin B. Importantly, cyclin B overexpression in TEM4-depleted cells largely rescues mitotic progression and chromosome segregation defects in anaphase. Our study thus illustrates the consequences of Rho signalling imbalance on cell cycle progression and identifies TEM4 as the first GEF governing Rho GTPase-mediated regulation of G1/S.

## Introduction

The eukaryotic cell division cycle is a complex process that necessitates remodelling of numerous architectural features of the cell including the microtubule and actin cytoskeletons. It is well-documented for example that interfering with microtubule polymerization using drugs such as nocodazole, colchicine or vinca alkaloids prevents proper spindle formation, and results in arrest of cells in prometaphase due to spindle assembly checkpoint (SAC) activation (Jordan et al., 1992; McAinsh and Kops, 2023). The actin cytoskeleton on the other hand is a major integrator of the various inputs that control G1/S transition. These inputs include both soluble stimuli, such as growth factors, and mechanotransduction cues from cell adhesions and the local microenvironment and are long known to synergistically control cell cycle progression with Rho GTPases being shared effectors of these pathways (Assoian and Schwartz, 2001; Boonstra and Moes, 2005; Hall, 2005; Jones et al., 2019; Reshetnikova et al., 2000; Uroz et al., 2018).

Among the numerous activities driven by Rho family GTPases, changes in cell morphology, adhesion, motility, cell cycle progression, survival, apoptosis and cytokinesis are well described (Hall, 2012; Jaffe and Hall, 2005; Lawson and Ridley, 2018). Through fine-tuned temporal and spatial regulation of the Rho GTPases, guanine nucleotide exchange factors (GEFs) contribute to the regulation of both actin and microtubule cytoskeletons (Rossman et al., 2005; Zuo et al., 2014). For example, recruitment of the RhoGEF Ect2 to the central spindle is essential for cytokinesis in most animal cells and inhibition of this localization prevents accumulation of RhoA, F-actin, phospho-myosin light chain, and anillin at the cortical membrane adjacent to the central spindle, all of which are necessary for initiation and ingression of the cleavage furrow (Nishimura and Yonemura, 2006; Somers and Saint, 2003; Su et al., 2011; Yüce et al., 2005). In addition to Ect2, several other GEFs have been implicated in cytokinesis including GEF-H1 (Birkenfeld et al., 2007), Myo-GEF (Wu et al., 2006) and LARG (Martz et al., 2013). In early mitosis, Ect2 regulates cortical rigidity of the plasma membrane. Upon mitotic entry, Rho A activity drives cortical actin filaments to form a meshwork at the cell surface that is exquisitely tuned to achieve optimal levels of cortex thickness and tension, and both too much or too little Rho A activity can deregulate this process (Chugh et al., 2017; Maddox and Burridge, 2003). The tensile properties of the cortex ultimately result in the characteristic rounded-shape of mitotic cells, a critical feature of cell division. This process is overseen by CDK1 activity as cells begin to enter mitosis, and numerous CDK1 targets have been implicated in this process (Chen et al., 2022; Jones et al., 2019, 2018; Nishimura et al., 2019; Watanabe et al., 1999).

In interphase, Rho A activity is important for passage through G1. Early microinjection studies using dominant-negative constructs or constitutively active mutants of Rho A, Rac1, cdc42, or the Rho toxin C3 transferase demonstrated the contribution of these to mitogen-induced G1 progression (Olson et al., 1995; Yamamoto et al., 1993). The major targets of Rho GTPase signaling within the cell cycle machinery are the CKIs p21^waf1/cip1^ (p21) and p27^kip1^. Dominant-negative Rho A was shown to increase p21 levels and, conversely, activated Rho A mutants prevented its up-regulation (Adnane et al., 1998; Olson et al., 1998). Rho inhibits p21 expression at the transcriptional and post-transcriptional level, and this inhibition may be essential for the effects of Rho on cell cycle progression (Adnane et al., 1998; Coleman et al., 2006; Han et al., 2005; Liberto et al., 2002; Olson et al., 1998; Song et al., 2001). The upstream GEFs that regulate Rho A in this context remain unknown.

ARHGEF17 (also known as and subsequently referred to as Tumor Endothelial Marker 4, TEM4) is an understudied GEF with three annotated domains: an N-terminal actin binding domain (ABD), a central catalytic DH-PH domain, and a C-terminal WD40 fold (Fig. 1A). TEM4 binds specifically and directly to dynamic, newly assembled F-actin filaments, via the ABD (Mitin et al., 2012; Prifti et al., 2022). In vitro studies indicated that the catalytic activity of TEM4 is specific for Rho over Rac1 and Cdc42 (Bagci et al., 2020; De Toledo et al., 2000; Mitin et al., 2012; Rümenapp et al., 2002a). Measurement of TEM4 activity in cells showed that binding to actin, which is required for its subcellular localization, may directly regulate TEM4 activity, as a mutation that abolished actin binding decreased TEM4’s capacity to activate Rho (Mitin et al., 2012). In addition, TEM4 has also been implicated in the maintenance of cell-cell adhesion, and in the formation of cell junctions and endothelial barriers in Madin–Darby canine kidney (MDCK) cells (Ngok et al., 2013). In agreement with the original identification of TEM4 as a tumor endothelial marker, García-Jiménez et al., proposed that TEM4 may be implicated in tumor growth and metastatic dissemination of lung cancer cells (García-Jiménez et al., 2022). Overall, the emergent picture suggests that TEM4 functions as a RhoGEF specifically activated by dynamic changes in the actin cytoskeleton to regulate functions related to cell shape, movement and contractility. In addition to these, the Mitocheck consortium (Neumann et al., 2010) revealed a cell cycle function for TEM4 and in a follow-up study, Isokane et al., proposed that it may be a novel regulator of the SAC during mitosis (Isokane et al., 2016).

**Fig. 1.**
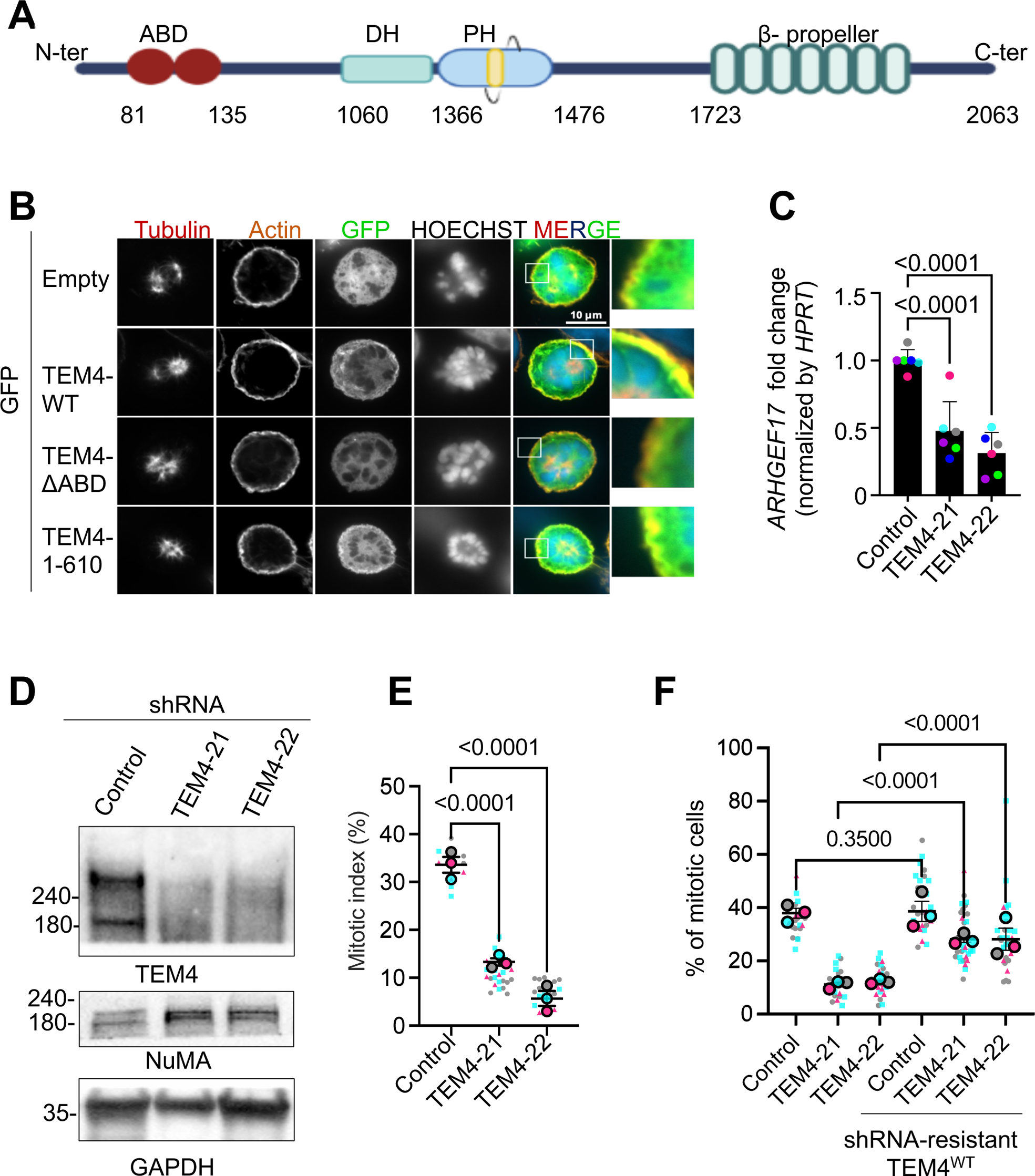
TEM4 is required for mitosis. A: Schematic illustration of the annotated domains of TEM4 protein. B: Representative immunofluorescence image of mitotic HeLa-T-REx cells overexpressing the indicated GFP-TEM4 constructs. Cells were stained with anti-GFP (green), Phalloidin-Atto 565 for F-actin (red) and Hoechst (blue). Scale bar = 10μm. C: Fold change in *ARHGEF17* mRNA in the indicated cell lines after 72h shRNA induction. Gene expression was normalized to HPRT. Shown is mean±SEM, n=6. D: TEM4 Western blot of lysates from the indicated cell lines after synchronization in mitosis with 100ng/ml nocodazole at the end of a 72h shRNA induction; n=3. E: The mitotic index of HeLa-T-REx control, TEM4-21 and TEM4-22 cells 72h after shRNA induction. Cells were treated with 100ng/ml nocodazole for 12 h prior to fixation. Data shown is mean ±SEM; n=3, N≥1000 cells. F: Control TEM4 shRNA and rescue cell lines were treated as indicated in E. Mean mitotic index ± SEM is show, n=3.

In the present work we investigate the role of TEM4 in controlling mitotic events. We find that TEM4 plays a crucial role in cell cycle progression with loss of TEM4 resulting in a significant block during G1 and delayed entry into mitosis. In the small proportion of cells that do enter mitosis, we identify defects in mitotic rounding, the timing of G2/M transition, and the duration of mitosis, as well as defects in spindle orientation and chromosome alignment and segregation. Mechanistically, we found that loss of TEM4 resulted in decreased transcription of *CCNB1* and thus reduced cyclin B protein, likely as an indirect consequence of increased levels of the G1 inhibitor p21 and a delayed cell cycle. Expression of exogenous cyclin B in TEM4-depleted cells, restored progression through mitosis, reinstated normal actin levels at the cortex and rescued chromosome segregation defects observed in the absence of TEM4. These observations collectively support the conclusion that TEM4 regulation of the actin cytoskeleton in interphase drives a transcriptional programme that promotes timely cell cycle progression. To the best of our knowledge, TEM4 is the first Rho regulator implicated in this process.

## RESULTS

### TEM4 is required for entry into mitosis

We first sought to determine TEM4 localization during mitosis. To do this, we expressed GFP-tagged full length TEM4 (GFP-TEM4^WT^), TEM4 lacking 11 amino acids from the ABD (residues 125-135) (GFP-TEM4^ΔABD^) and a construct entirely lacking the C-terminus of TEM4 (GFP-TEM4^1-610^) (Mitin et al., 2012). In agreement with Mitin et al., we confirmed that in interphase, full length TEM4 localizes at the actin cytoskeleton and that the ABD is required for this localization, as deletion of this domain resulted in diffuse cytoplasmic staining and loss of colocalization with actin stress fibers (Mitin et al., 2012) (Fig. S1). During mitosis, TEM4 mostly localized to the actin cortex in a manner similarly dependent on the ABD, but in contrast to the kinetochore localization that has been previously reported (Fig. 1B)(Isokane et al., 2016). To study the function of TEM4 in mitosis, we first generated inducible HeLa-TRex cell lines expressing two independently verified shRNAs targeting TEM4 (hereafter referred to as TEM4-21 and TEM4-22) (Isokane et al., 2016; Memon et al., 2021; Ngok et al., 2013; Prifti et al., 2022; Weber et al., 2021). We initially validated the efficiency of TEM4 depletion by qPCR and detected a 50-60 % decrease in TEM4 mRNA levels 72 hours post shRNA induction (Fig. 1C). Using an in-house generated antibody, Western blotting of mitotic cell extracts showed a clear decrease in the protein levels of TEM4 after induction of both shRNAs, (Fig. 1D, see materials and methods for antibody details). To evaluate the effect of TEM4 depletion on mitotic progression, we measured the mitotic index in cells depleted of TEM4 and synchronized in mitosis using nocodazole. We found a clear decrease of mitotic cells in the absence of TEM4 (Fig. 1E) as has been previously reported (Isokane et al., 2016; Memon et al., 2021; Weber et al., 2021). The low mitotic index was observed also in the presence of taxol, and in cells synchronized in mitosis by release from a double thymidine block, suggesting that the delay is independent of the status of microtubules and the method of synchronisation (Fig. S1B-C). Moreover, TEM4 depleted cells were unable to accumulate in mitosis even in the presence of the proteosome inhibitor MG132 indicating that the low mitotic index was not a result of accelerated cell cycle progression (Fig. S1D). Importantly, mitotic progression was largely restored in cell lines rescued with shRNA resistant TEM4^WT^ in both shRNA conditions (Fig. 1F, Fig. S1E, F) demonstrating specificity of knockdown. The low mitotic index observed in HeLa cells was reproducible in HCT-116 cells similarly engineered to express inducible depletion of TEM4 (Fig. S1G-J).

### TEM4 depletion delays mitotic entry and progression and results in multiple mitotic aberrations

To better understand the effect of TEM4 depletion on mitotic events we turned to live cell imaging. Cells were synchronized in G1 by thymidine then released and monitored for mitotic entry. Live cell imaging confirmed that loss of TEM4 had a profound effect on the ability of cells to both enter and progress through mitosis. Whereas control cells entered mitosis 8.7 ±1.4 h post release from thymidine, TEM4-21 cells required 12.5 ±3.15 h and TEM4-22 cells 13.6 ±3.2 h before entering mitosis (Fig 2A, Fig. S2A). Entry into mitosis was also remarkably asynchronous for TEM4-depleted cells relative to controls (Fig. 2A, S2B). In addition, mitotic duration was significantly prolonged in TEM4-depleted cells averaging 100±40 mins and 70±27 mins for TEM4-21 and TEM4-22, respectively, compared to 38 ±11.06 mins for control cells, with the delay spread across all mitotic stages (Fig. 2B). Moreover, in cells that did enter mitosis after TEM4 depletion, live cell imaging revealed a significant increase in mitotic cell death in both cell lines depleted for TEM4 either soon after nuclear envelope breakdown or after multiple attempts at metaphase alignment (Fig. 2C).

**Fig. 2.**
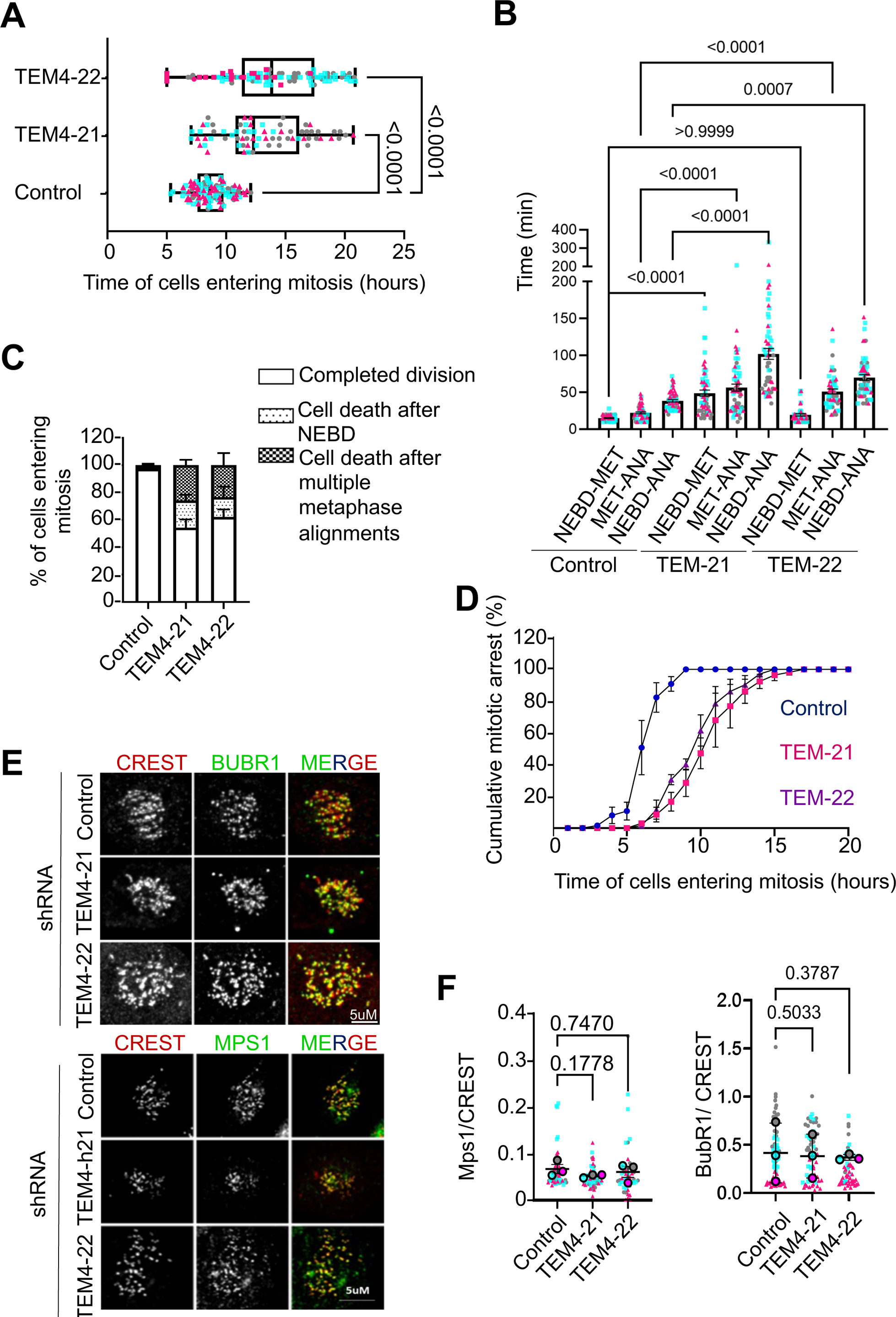
TEM4 depletion leads to mitotic delay, prolonged mitosis and cell death. A: Timing of mitotic entry after G1 release in the indicated cell lines. Individual data points are shown from 3 replicate experiments, N≥ 30 cells per condition. B: Timing of mitotic transitions from the cells in (A); NEBD: nuclear envelope breakdown, MET: metaphase, ANA: anaphase. C: Quantification of the cell fate phenotypes from the live cell imaging experiment in (A). n=3, N≥ 200. D: Cumulative mitotic index in in nocodazole-treated HeLa -TREx cells expressing TEM4 shRNA after 72 h of induction as determined from live cell imaging. The graph displays the mean ± SD, n=3; N≥20 per condition. E: Control, TEM4-21, TEM4-22 cells were induced for 72h and 0.4 μΜ RO-3306 was added for the final 16h, before release into fresh media for 30 mins. Cells were stained with anti-CREST (red), Hoechst (blue), and either anti-MPS1 or anti-BUBR1 (green) as shown. Scale bar = 5μm. F: Quantification of average relative intensity of MPS1 and BUBR1 at kinetochores from (E). Graph shows mean ± SEM; n=3, N≥20 cells/ condition/ experiment.

TEM4 was identified as an essential mitotic gene by the Mitocheck consortium and has been implicated in SAC activation through ensuring timely localization of MPS1 at kinetochores (Isokane et al., 2016). However, our results so far indicate that TEM4 depletion leads to prolonged mitosis and delay of entry in the proportion of cells that do enter M-phase. In order to better address these discrepancies, we tested directly the ability of TEM4 depleted cells to arrest in the presence of nocodazole by live-cell imaging. We found that although TEM4 depleted cells entered mitosis sparingly and relatively late as shown above in the presence of nocodazole (Fig. 2A, S2B), TEM4 depletion did not result in premature exit from mitosis with cells remaining arrested in prometaphase for the duration of filming (Fig. 2D). As an additional validation of spindle checkpoint function, we monitored kinetochore localization of MPS1 and Budding Uninhibited by Benzimidazole 1-Related 1 (BUBR1). We found that TEM4 depletion did not significantly reduce MPS1 or BUBR1 at the kinetochore, suggesting that for the small percentage of cells that do enter mitosis in the absence of TEM4, loss of SAC signaling was not a major consequence (Fig. 2E, F).

### TEM4 depletion leads to aberrant spindle formation, orientation and chromosome segregation defects

The cortical localization of TEM4 and delayed mitotic progression in TEM4-depleted cells led us to postulate that TEM4 may have a role at the mitotic cortex. To test this idea, we quantified spindle phenotypes in our live cell imaging which revealed significant abnormalities including spindle rotation defects, multiple rounds of spindle metaphase alignment, and cytokinesis defects (Fig. 3A, see also supplemental movies Video1, 2,3). These observations were corroborated by measurement of spindle angles in fixed cells which revealed that compared to control cells, the majority of cells depleted of TEM4 with either shRNA exhibited spindle angles between 40 and 90 degrees, while the majority of control cells had angles between 0 and 30 degrees relative to the coverslip, indicating increased spindle movement away from the plane of division in TEM4 depleted cells (Fig. 3B).

**Figure 3:**
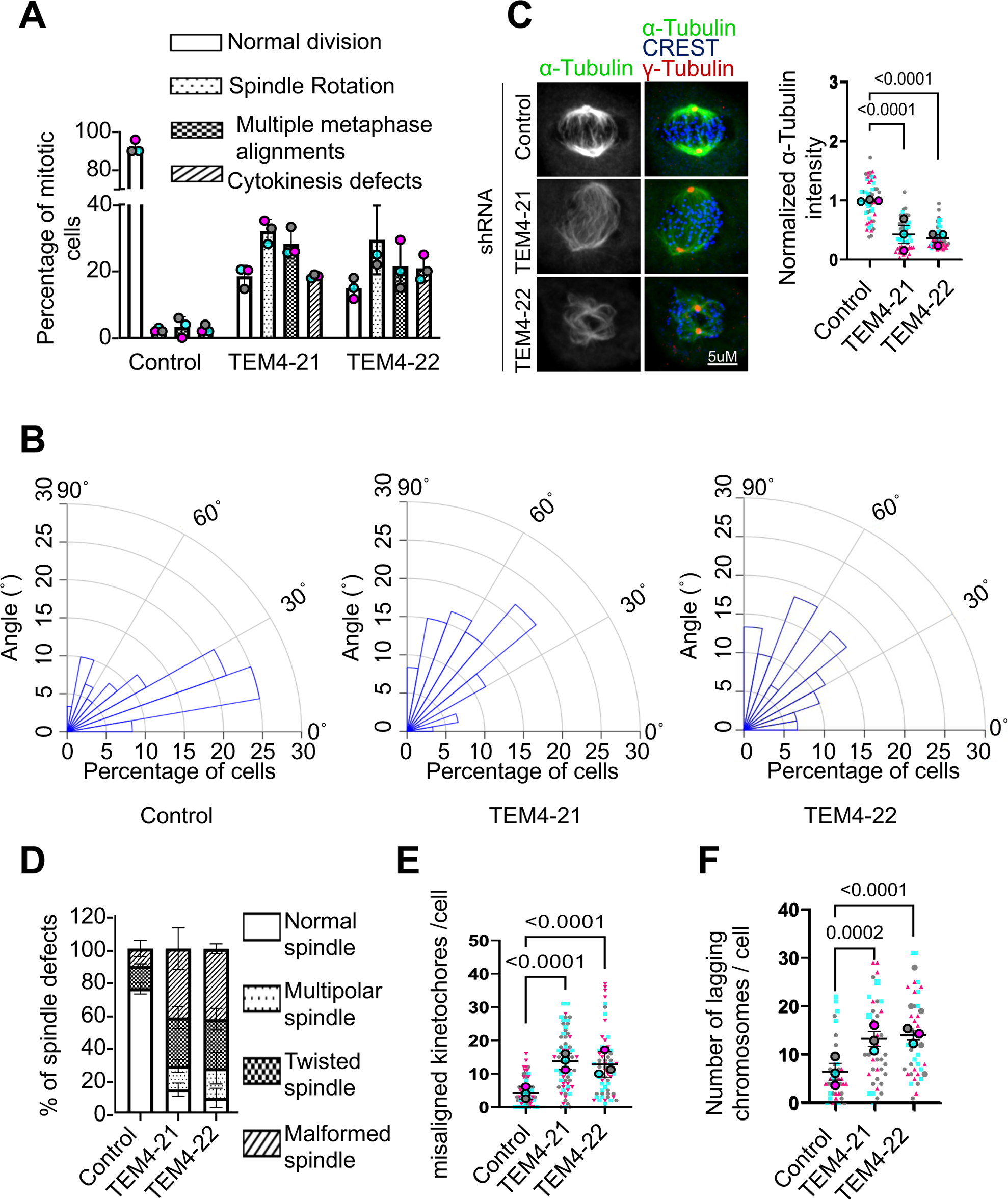
TEM4 depletion leads to errors in spindle formation and orientation and chromosome segregation defects. A: Quantification of the most common spindle phenotypes from Fig. 2A. A total of approximately 200 cells were analyzed from 3 independent experiments. B: Spindle angle calculations from control, TEM4-21 and TEM4-22 cells. n=3, N≥20 cells/condition. C: ShRNA expression was induced for 72 h using 0.5 μg/ml doxycycline and cells treated for 3 h with 3μM of nocodazole. After nocodazole washout cells were incubated with fresh media for 2 h. Fixed cells were stained with CREST (blue), anti-α-tubulin (green) and anti-γ-tubulin (red). Mean± SEM α-tubulin intensity 2 h post release is shown, n=3, N≥20 cells/condition). Scale bar = 5μm. D: Quantification of the spindle phenotypes observed in C. E: Quantification of chromosome misalignment in parental, TEM4-21 and TEM4-22 cells after shRNA induction. Cells were treated for 12 h with 5 μM STLC and released into fresh media containing MG132 for 4 h, prior to fixation and staining. Large symbols represent the means ± SEM of individual experiments. Statistical differences were measured by a repeated-measures two-way ANOVA on the average of each condition for each of 3 biological replicates (≥20 cells/condition). F: Quantification of lagging chromosomes in anaphase. Cells were treated for 12 h with 0.4 μΜ RO-3306 and released into fresh media for 1.5 h for control cells and 2.5 h for TEM4 shRNA inducible cell lines to enrich for cells in anaphase. Data shows means ± SEM, n=3, N≥25 cells/condition.

We next examined spindle formation in the absence of TEM4 by monitoring microtubule regrowth after nocodazole induced depolymerization. As expected after nocodazole washout, microtubules repolymerized in a radial manner, forming a distinct aster-like structure and bipolar spindles within an hour in control cells. In contrast, depletion of TEM4 resulted in slower microtubule growth and the formation of multipolar spindles (Fig. S3A), in agreement with our observations by live cell imaging discussed above (Fig. 3A). Furthermore, even two hours after release from nocodazole, TEM4 depleted cells exhibited reduced microtubule density consistent with decreased spindle reformation capacity (Fig. 3C). Quantification of the spindle defects at the same time point revealed that whereas 75.9 ± 3.6% of control cells displayed normal mitotic spindles, only 14.0± 4.0% of TEM4-21 and 8.97± 5.9% of TEM4-22 expressing cells were able to form normal spindles. For both cell lines depleted of TEM4, the remaining cells displayed multiple spindle defects including multipolar and twisted spindles (Fig. 3D. Fig. S3B). A major consequence of defective spindles is the misalignement of chromosomes at metaphase and their missegregation at anaphase. To test for alignment, we treated control and TEM4 depleted cells with the Eg5 inhibitor S-Trityl-L-Cysteine (STLC) to disrupt the formation of bipolar spindles. Cells were then released into media containing the proteasome inhibitor MG132 to allow for bipolar spindle formation and alignment, and kinetochore positions relative to the metaphase plate were monitored as previously reported (Gama Braga et al., 2021). We found that cells depleted of TEM4 exhibited a significant increase in misaligned kinetochores compared to control cells (Fig. 3E). In cells that were allowed to progress to anaphase in the absence of MG132, TEM4 depletion resulted in a marked increase in lagging chromosomes (Fig. 3F). Taken together, our data suggest that TEM4 expression is crucial for proper spindle formation and chromosome segregation during mitosis.

### TEM4 depletion leads to Rho A activation resulting in excessive cortical F-actin and loss of mitotic retraction fibres

At mitotic entry, reorganization of the actin cytoskeleton is driven by Rho A and results in cortical stiffness and cell rounding (Maddox and Burridge, 2003; Matzke et al., 2001). Early studies also demonstrated that F-actin coordinates cell margin retraction during rounding, and that this process is dependent on tightly regulated Rho A activity (Chugh et al., 2017; Cramer and Mitchison, 1997; Maddox and Burridge, 2003). In addition, actin-rich retraction fibers, which maintain the connection of rounded mitotic cells to the adhesive substrate, contribute to correct spindle orientation parallel to the substratum and ensure attachment of both daughter cells after division (Théry et al., 2007; Toyoshima and Nishida, 2007). Given the multiple spindle defects we observed in TEM4 depleted cells (Fig. 2A-D) as well as its cortical localization, we asked if TEM4 depletion could contribute to actin cortex functions in metaphase cells. We found that whereas control cells or non-induced cells expressing TEM4 shRNA exhibited clear retraction fibres, 72h after TEM4 shRNA induction, the percentage of cells exhibiting clear retraction fibres were significantly reduced in TEM4 depleted cells (Fig. 4A, B). Moreover, and in agreement with observations that retraction fibres play an important role in maintaining the shape and roundness of mitotic cells (Cramer and Mitchison, 1997; Taubenberger et al., 2020), we found that TEM4 depletion resulted in a decrease in the circularity of mitotic cells (Fig. 4 C).

**Figure 4:**
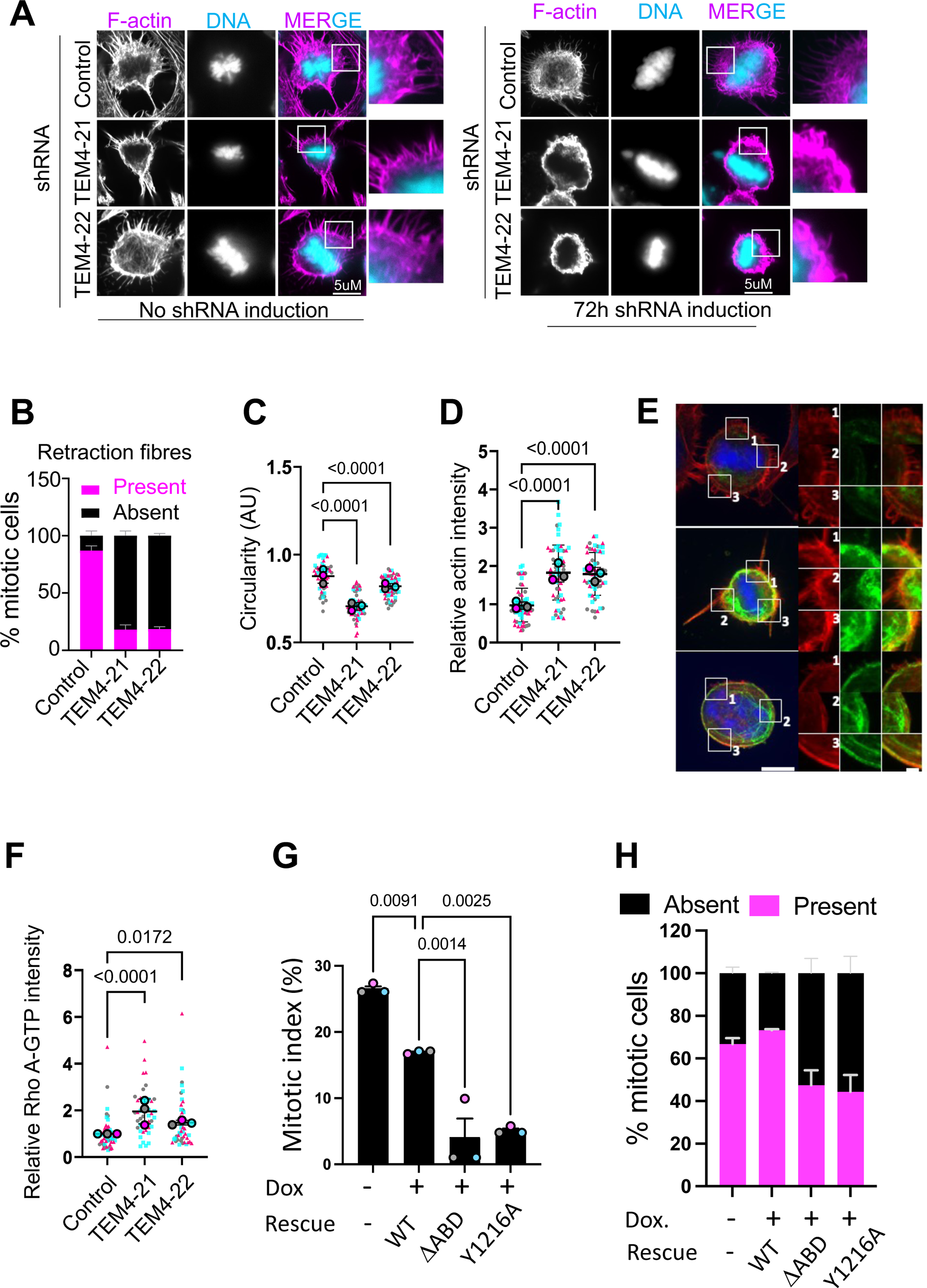
TEM4 depletion leads to excessive cortical actin and loss of retraction fibres in mitosis. A: Control, TEM4-21 and TEM4-22 expressing cells before and after 72h of induction were synchronized in mitosis after release from double thymidine block, then fixed before stained with Phalloidin-Atto 565 (magenta) and Hoechst (cyan). Scale bar = 5μm. B: The percentage of cells with or without retraction fibers was quantified in metaphase after 72h of TEM4 depletion. Mean percentages ± SEM is shown, n=3, N ≥20 cells/condition. C: Circularity in control and TEM4-depleted metaphase cells, where 1 is considered a perfect circle. Data shown is mean ± SEM with ≥20 cells/condition. D: Actin intensity at the cortex in control and TEM4-depleted metaphase cells, mean ± SEM (≥20 cells/condition). E: Immunofluorescence of Rho A-GTP (green) and F-actin (red) at the cell cortex of control and TEM4-depleted cells. Large scale bar =10μM, small scale bar =2μM F: Quantification of the experiment in E. Rho A-GTP at the cortex was measured in maximum projections of cells stained for active Rho A. Data shows mean ± SEM, n=3, N ≥20 cells/condition. G: Mitotic index in control cells, and cells rescued with TEM4^WT^, TEM4^ϕλABD^, TEM4^Y1216A^. Data shows mean ± SEM, n=3, N ∼100 cells per condition. H: Percentage of control cells, and cells rescued with TEM4^WT^, TEM4^ϕλABD^, TEM4^Y1216A^ exhibiting retraction fibres. Data shown is mean ± SEM, n=3, N ≥100 cells per condition.

In addition to the phenotypes described above, we observed a thickening of the cortex with increased F-actin levels in TEM4 depleted cells indicative of increased cortical contractility (Fig. 4 A, D). This increase in cortical actin was reproducible in additional TEM4 shRNA clonal lines indicating that this was not a clonal effect (data not shown). These results were surprising given that role of TEM4 as a Rho activator, and how loss of TEM4 could contribute to increased F-actin polymerization is considered in the discussion. Nevertheless, and to explore this phenotype in more detail, we first used polarization microscopy to confirm decreased cell contractility in interphase cells depleted of TEM4 (Fig. S4A). The results are in agreement with role for TEM4 in Rho A activition (Lutz et al., 2013; Ngok et al., 2013; Rümenapp et al., 2002b). In addition, we used a Rho-GTP-specific antibody to observe active RhoA localization in TEM4-depleted cells. Although, as expected given the localization of TEM4, we did not observe obvious global effects on Rho A activation, we noted decreased levels of active Rho A in lamellipodia and other F-actin rich structures in TEM4-depleted cells relative to controls (Fig. S4B). These results indicate that in TEM4-depleted cells, Rho A activity is attenuated. Although technical limitations prevented the use of fluorescence polarization in mitotic cells, visualization of RhoA-GTP at the cortex by immunofluorescence demonstrated a small but consistent increase in active Rho A in TEM4 depleted cells, in agreement with the increased F-actin density and contractility reported above (Fig. 4E, F). As a further readout of Rho A activity, we also measured by immunofluorescence phosphorylation of the ERM proteins, known regulators of cortical contractility, and spindle organization and positioning (Carreno et al., 2008; Hirao et al., 1996; Kunda et al., 2008) and found increased phosphorylation in TEM4-depleted cells compared to control cells consistent with Rho A hyperactivation (Fig. S4C).

To determine whether the cortical phenotypes in mitosis were dependent on TEM4 GEF activity or cortical localization, we generated additional stable cell lines expressing GFP-3xMYC-tagged TEM4^ΔABD^, and the GEF-inactiveTEM4^Y1216A^ in each of the TEM4-21 and TEM4-22 backgrounds (Fig. S5) and determined the capacity of these to rescue mitotic phenotypes relative to GFP-3xMYC-TEM4 WT. We found that whereas TEM4^WT^ partially rescued the mitotic index and retraction fibres formation, TEM4^Y1216A^ and TEM4^ΔABD^ were unable to do so (Fig. 4F, E). Taken together, our results demonstrate a requirement for TEM4 GEF activity and actin binding capacity for mitotic entry and for the proper regulation of F-actin at the cell cortex.

### Rho kinase inhibition rescues mitotic actin contractility and cell cycle progression in TEM4 depleted cells

The cortical actin network that is assembled in early mitosis is dependent on Rho A regulation of the formin DIAPH1 that drives cortical assembly of a non-branched actin network and the Rho kinases (ROCK1 and ROCK2) that activate non-muscle myosin II to crosslink the actin filaments generated by DIAPH1 thereby increasing contractility and tension (Lancaster and Baum, 2014; Rizzelli et al., 2020; Taubenberger et al., 2020). Given the observed Rho A hyperactivation, and increased contractility, we reasoned that inhibition of ROCK should rescue the associated phenotypes. Cells were therefore released from a double thymidine block and monitored for mitotic entry at which point they were treated for 30 minutes with the ROCK inhibitor Y-27632, (Ishizaki et al., 2000). This inhibition resulted in an increase in the percentage of TEM4 depleted cells with retraction fibers but had little effect on control cells (Fig. 5A, B). Additionally, actin intensity at the cortex decreased significantly in TEM4 depleted cells treated with Y-27632 while it remained unchanged in control cells (Fig. 5C). Moreover, we found that ROCK inhibition upon mitotic entry slightly but significantly increased the percentage of cells in mitosis under conditions of TEM4 depletion, indicating rescue of the slow mitotic passage (Fig. 5D). Overall, our data show that the effects of increased Rho A activation in TEM4 depleted cells on the cortex in mitosis can be restored, at least in part, by inhibition of downstream ROCK signalling.

**Figure 5:**
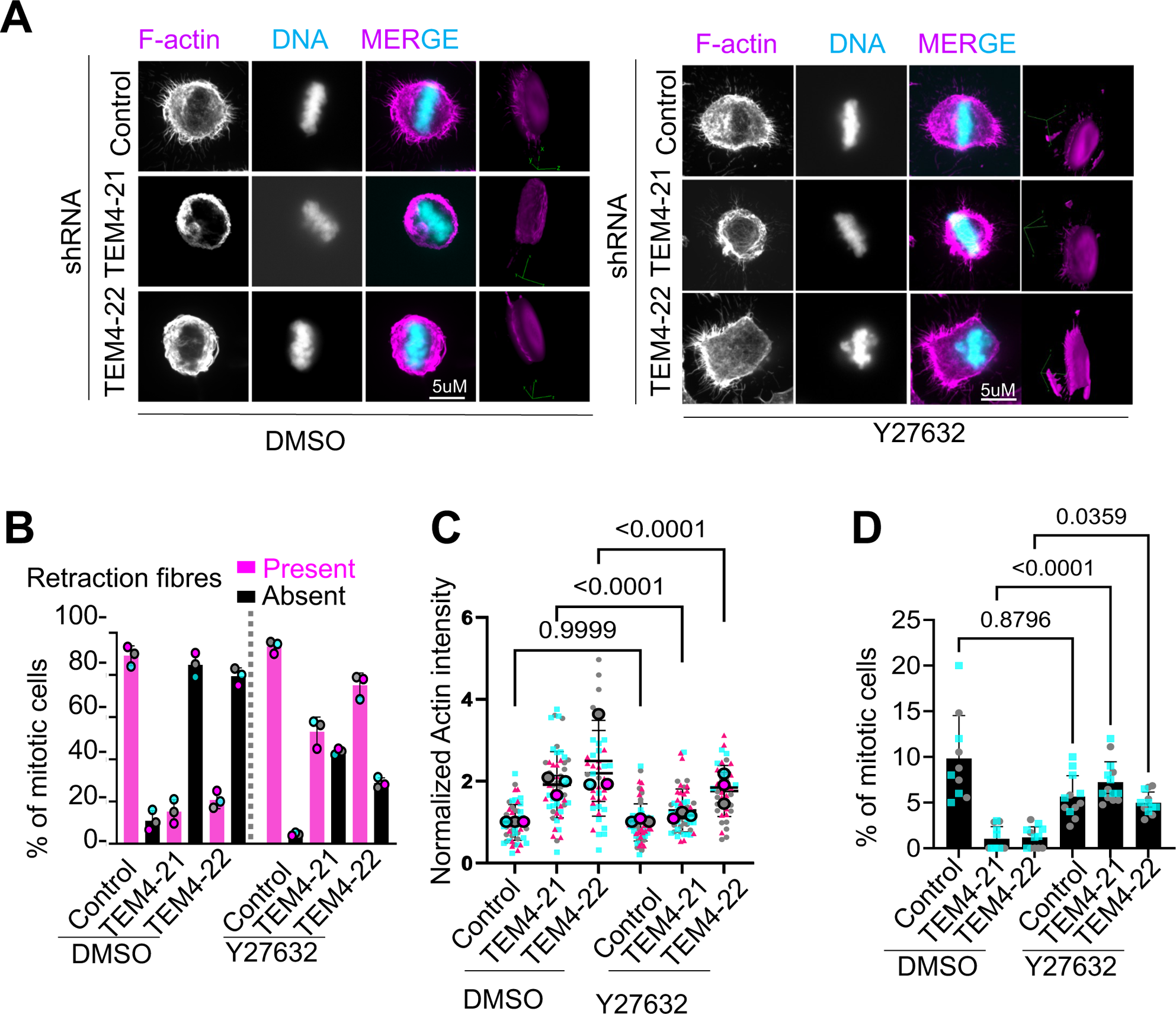
Rho kinase inhibition restores retraction fibres and reduces accumulation of f-actin at the cortex. A: Control and TEM4-depleted cells were synchronized in mitosis after release from a double thymidine block and treated for 30 mins with DMSO or the ROCK inhibitor, Y27632 prior to fixation. Cells were stained with Phalloidin-Atto 565 (magenta) and Hoechst (cyan). Scale bar = 5μm. Orthogonal views are shown in the rightmost panel. B: The percentage of cells from (A) with or without retraction fibers was quantified in metaphase cells in the presence of DMSO or Y27632. Data show is mean ± SEM, n=3, N ≥50 cells/condition. C: Quantification of cortical actin intensity of cells from (A). Data shown is mean ± SEM, n=3,N ≥20 cells/condition. D: Mitotic index in cells treated as in A. Data is shown from two representative experiments with ≥100 cells/condition. Individual datapoint represent individual fields of imaging within an experiment.

### TEM4 depletion results in G1 delay and decreased expression of the mitotic cyclins

Thus far, our data are generally consistent with a model whereby TEM4 acts as a cortex-bound Rho exchange factor that coordinates proper formation of retraction fibres and cell contractility in early mitosis. To provide further evidence for this, we sought to demonstrate that the delay in mitotic entry resulted in an accumulation of cells in G2. We treated cells with nocodazole for 12 h and determined the cell cycle profile in control and both TEM4 shRNA cell lines using FACS. To our surprise, we found that while more than 60% of control cells accumulated in G2/M under these conditions, TEM4 depleted cells remained mostly in G1 with less than 30% progressing to G2/M (Fig.6A). In an effort to understand the mechanisms through which TEM4 regulates cell cycle progression, we turned initially to recent phosphoproteomic analysis performed in cancer cells depleted of endogenous TEM4. Using an integrated phosphoproteomic, genomic and transcriptomic approach, TEM4 was identified as a candidate gene likely to drive changes in kinase signalling in the Hippo pathway (Memon et al., 2021). As part of this work, Memon *et al*. generated phosphoproteomics data from T47D and MDA-MB-468 breast cancer cell lines depleted of TEM4. Notably, these authors used TEM4 shRNA duplexes largely overlapping TEM4-22. Gene ontology (GO) enrichment analysis of phosphosites downregulated (log_10_-fold<-0.5) in TEM4-depleted cells from these datasets revealed that in addition to changes in phosphorylation of proteins associated with “actin cytoskeleton organization” as anticipated for a RhoGEF, we observed enrichment of terms related to the progression of the cell cycle including “regulation of cell cycle processes”, “microtubule cytoskeleton organization” and “mitotic cell cycle process” (Fig. S6A, B, Supplemental table 1). Importantly the enrichment of these terms was observed in both T4D and MDA-MB-468 cell lines only in the sites that were hypophosphorylated and was not observed for hyperphosphorylated sites (not shown). Next, we identified all downregulated phosphosites in TEM4 depleted cells in proteins with GO terms associations related to cell cycle progression and mitosis. In MDA-MB-468 cells, this led to the identification of 152 unique phosphosites in 111 proteins of which 43 (∼28%) were high confidence phosphosites (probability of phosphosite assignment >94%) corresponding to a minimal CDK motif (SP/TP). In T47D cells, we found 85 unique phosphosites in 69 proteins of which 32 (∼38%) were high confidence sites matching the minimal CDK target motif (Supplemental table 2). These data suggest that loss of cyclin-CDK phosphorylation is prevalent in cells depleted of TEM4.

**Figure 6:**
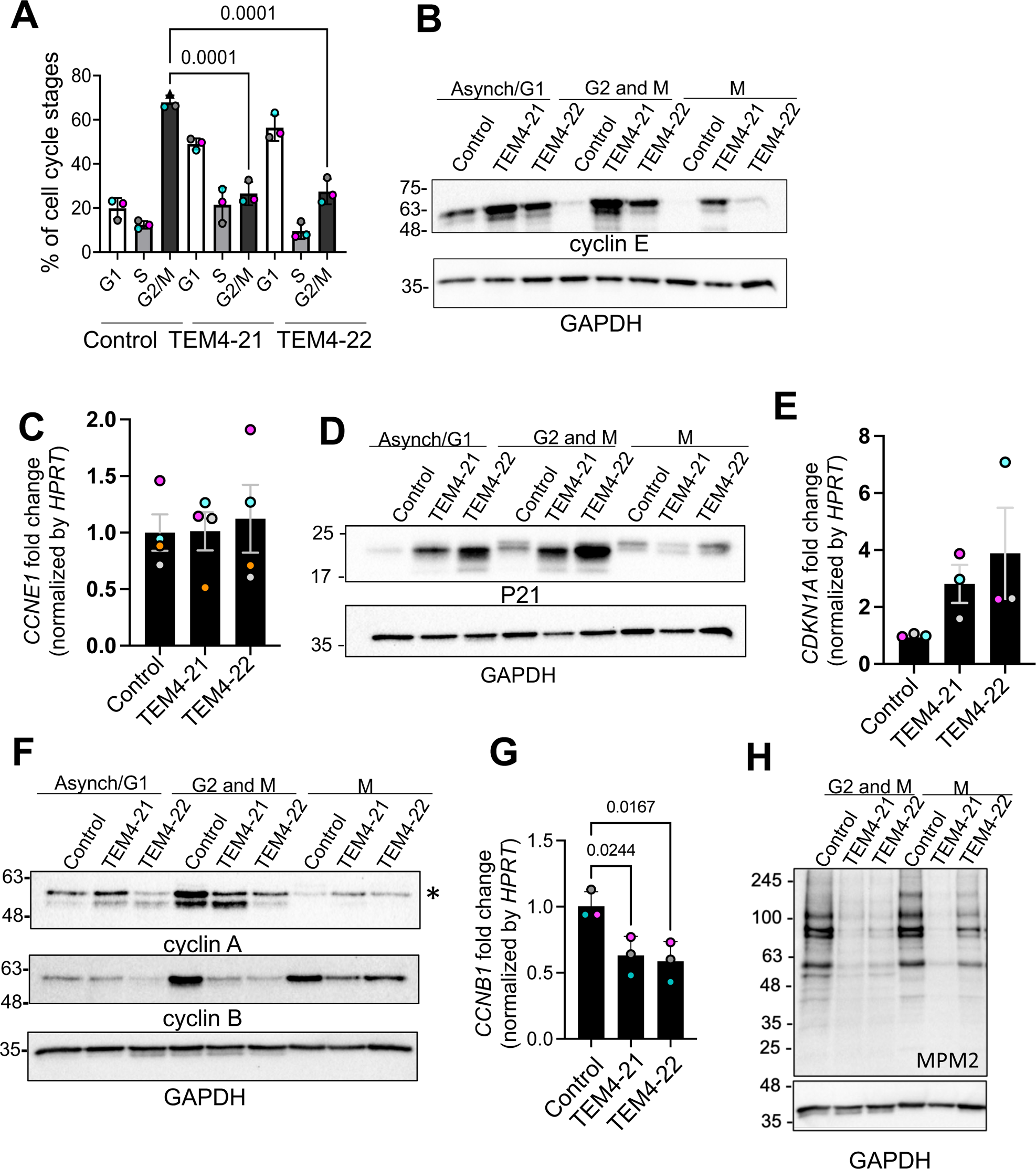
TEM4 depletion results in G1 delay and decreased expression of the mitotic cyclins. A: Cell cycle distribution of control and TEM4-depleted cells measured by FACS. The indicated cell lines were treated for 12 h with 100ng/ml nocodazole before being fixed and stained with Hoechst. Data shown is mean ± SEM, n=3. B: Control, TEM4-21, and TEM4-22 cells were left asynchronous, synchronized at G2/M with nocodazole or in mitosis after a nocodazole arrest and mechanical dislocation of round mitotic cells. Equalized lysates were then blotted for the indicated antibodies. C: Expression of *CCNE1* in interphase was measured by qPCR the indicated cell lines and expressed as fold change relative to *HPRT.* Data shows mean ± SEM, n=4. D: Control, TEM4-21, and TEM4-22 cells were treated as in (B) and lysates were probed with the indicated antibodies. E: Expression of *CDKN1A* in interphase was measured by qPCR the indicated cell lines and expressed as fold change relative to *HPRT.* Data shows mean ± SEM, n=3. F: Control, TEM4-21, and TEM4-22 cells were treated as in (B) and lysates were probed with the indicated antibodies. G: Expression of *CCNB1* in interphase was measured by qPCR the indicated cell lines and expressed as fold change relative to *HPRT.* Data shows mean ± SEM, n=3. H: Lysates from cells synchronized in G2/M and M phase as indicated in (B) and blotted for the indicated antibodies.

To determine the mechanism through which TEM4 regulates cell cycle progression, we first sought to explore the effect of TEM4 on the protein and mRNA levels of cyclin proteins and their inhibitors. For this, lysates were prepared from three populations of control and TEM4-depleted cells (enriched in G1, enriched in G2-M and M-phase cells, see matherials and methods for synchronization details). In both TEM4 shRNA cell lines, we found increased levels of cyclin E protein but not mRNA compared to control cells in all even in mitotic cells generated by mechanical “shakeoff”, suggesting that TEM4 depletion impeded efficient degradation of cyclin E (Fig. 6B, C), and in agreement with the delay of these cells in G1/S. Moreover, we found TEM4 depleted cells resulted in elevated protein levels of the cyclin E/CDK2 inhibitor p21 in both interphase and G2-M cells, as well as elevated mRNA levels in interphase cells compared to control (Fig. 6D, E). These observations strongly suggest that TEM4 depletion slows cell cycle progression via upregulation of the cyclin E-CDK2 inhibitor p21 and explain the decrease in CDK target phosphorylation despite increased levels of cyclin E.

We next determined the effect of TEM4 depletion on the expression and activity of the late cyclins, cyclin A and cyclin B. Compared to control, we consistently observed a reduction in cyclin B and cyclin A protein in both G2-M and M populations in TEM4 depleted cells (Fig. 6F). By qPCR, we found *CCNB1* transcripts from mitotic cells were significantly lower than in control cells (Fig. 6G). Finally, we show that global Ser/Thr phosphorylation (as measured by the anti-pSer/pThr-Pro MPM2 antibody) is severely reduced in G2-M and M lysates compared to controls indicating reduced CDK activity. Overall, these observations explain the G1 delay seen in TEM4 depleted cells as well as the breadth of phenotypes seen in TEM4-depleted mitotic cells and indicate that loss of TEM4 results in a loss of proper cyclin expression culminating in attenuated cyclin B - CDK1 activity in mitosis.

### Expression of exogenous cyclin B rescues cortical and mitotic phenotypes in TEM4 depleted cells

The observations above suggest that TEM4 may contribute to mitotic progression through ensuring proper expression of cyclin B. We therefore asked whether expression of exogenous cyclin B could rescue mitotic errors in TEM4 depleted cells. To this end, we transfected GFP-cyclin B into control, TEM4-21 and TEM4-22 cells (Fig. 7A), and first checked for rescue of actin-rich retraction fibres, a structure dependent on cyclin B-CDK1 activity (Chen et al., 2022; Jones et al., 2019, 2018; Robertson et al., 2015). In control cells overexpression of cyclin B1 had little effect on the percentage of cells with visible retraction fibres. In contrast, GFP-Cyclin B expression significantly increased the percentage of cells with retraction fibres in TEM4 depleted cells, indicating at least partial rescue of this phenotype (Fig. 7B). Next, we monitored the effect of GFP-cyclin B expression on the progression of mitosis. To do this, control and TEM4-depleted cells expressing GFP or GFP-Cyclin B were synchronized in mitosis after a release from thymidine before fixation and quantification of the distribution of mitotic stages in the different cell lines. While GFP-Cyclin B expression had little effect on the progress of control cells as illustrated by the relatively similar percentages of cells in the various mitotic stages in the presence or absence of exogenous cyclin B, and as previously reported (Jin et al., 1998; Laoukili et al., 2005), GFP-cyclin B expression shifted the profile of TEM4-21 and TEM4-22 cells relative to expression of GFP-alone, with a higher percentage of cells in anaphase and telophase in lieu of fewer cells in prometaphase (Fig. 7C). In agreement with these results, we find that the mitotic index in cells expressing GFP-cyclin B and treated with nocodazole was restored almost to control levels in TEM4-depleted cells (Fig. 7D). In addition, GFP-Cyclin B overexpression effectively rescued anaphase lagging chromosomes in TEM4 depleted cells, whereas overexpression of cyclin B increased lagging chromosomes in control cells as previously reported (Davydenko et al., 2013) (Fig. 7E). Taken together, our results suggest that the major consequences of the disrupted mitoses identified in TEM4 depleted cells can be rescued by restoring cyclin B.

**Figure 7:**
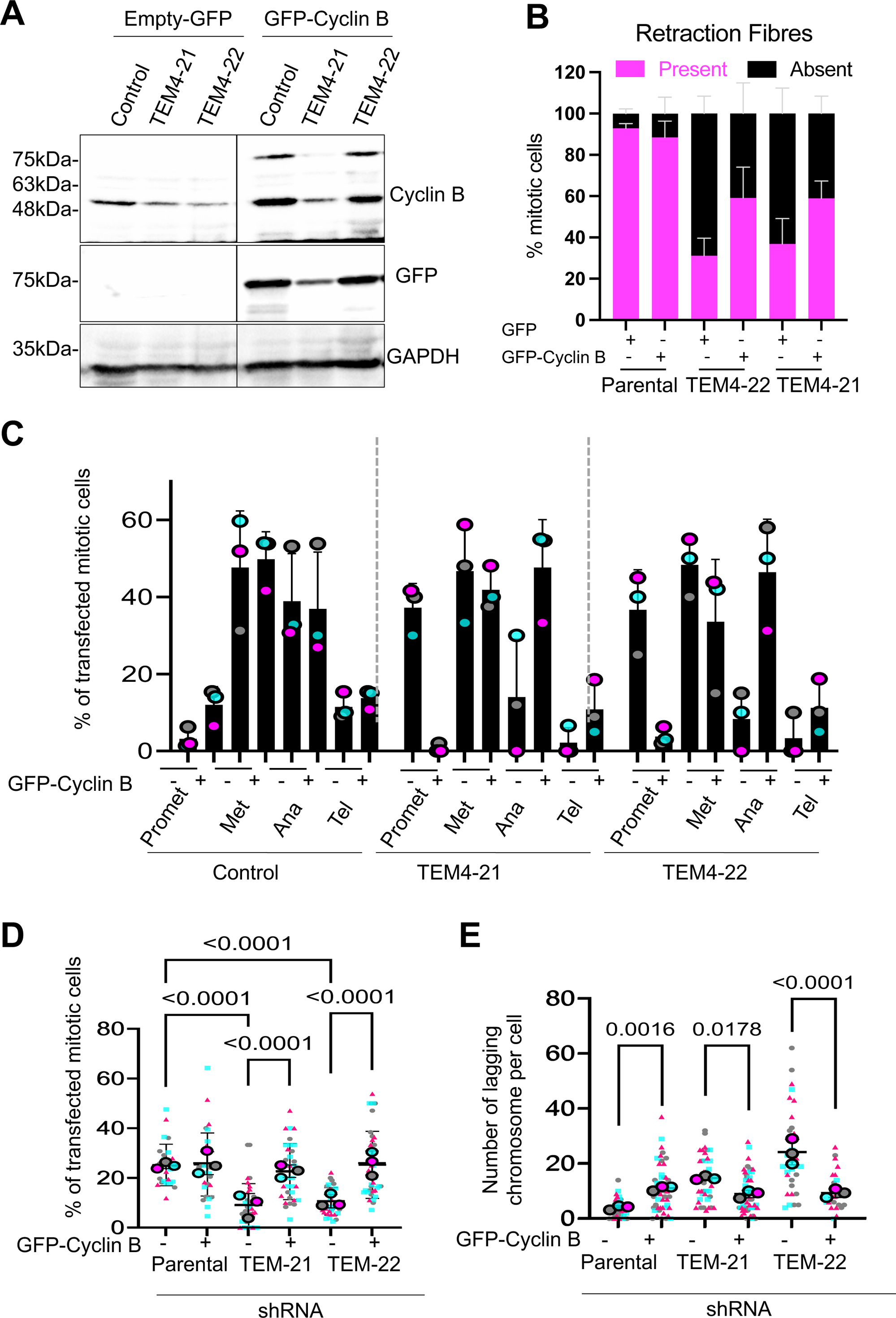
Overexpression of Cyclin B restores the mitotic index, cell cycle progression and reduces lagging chromosomes in TEM4 depleted cells. A: Control and TEM4-depleted cells were transfected with GFP- or GFP-cyclin B for 48 h prior total protein extraction with RIPA buffer. Western blotting was performed with the indicated antibodies. B: Cells grown on coverslips were transfected as in A and synchronized in mitosis with 100 ng/ml of nocodazole for 12 h prior fixation with PTEMF. Data shows mean ± SEM, n=3, N ∼200 cells. C: Mitotic progression of TEM4 depleted cells in the presence or absence of cyclin B. TEM4. Cells were transfected as in B and synchronized in mitosis after release from double thymidine block and the percentage of cells in each mitotic stage was calculated for all indicated conditions. Data shows mean ± SEM, n=3, N ≥ 50 cells per condition. D: Mitotic index in TEM4 depleted in the presence or absence of cyclin B. Cells were treated for 12 h with taxol. Data shows mean ± SEM, n=3, N≥25 cells per condition. E: Quantifications of lagging chromosomes in TEM4 depleted in the presence or absence of cyclin B. Cells were treated for 12 h with 0.4 μΜ RO-3306 and released into fresh media for 1.5 h. Data shows mean ± SEM, n=3, N ≥25 cells per condition.

## Discussion

TEM4 is an understudied RhoGEF implicated in numerous cell functions through its role in remodelling of the actin cytoskeleton. Here, we provide evidence that it promotes cell cycle progression at the G1/S transition, indirectly supporting efficient expression of mitotic cyclins and hence the accurate execution of cell division. Using two independently verified shRNAs throughout the study, we show that TEM4 depletion results in a marked delay in G1 despite high levels of cyclin E. This G1 delay is a result of elevated expression of the cyclin E-CDK2 inhibitor p21, levels of which have been shown increase upon Rho inhibition or depletion (Adnane et al., 1998; Coleman et al., 2006; Han et al., 2005; Liberto et al., 2002; Olson et al., 1998), and in cells treated with agents that prevent actin polymerization such as cytochalasin D and Latrunculin B (Coleman et al., 2006; Lee et al., 2009; Lohez et al., 2003). These data suggest that TEM4 regulation of Rho activity and downstream F-actin formation regulates p21 levels and reveal, to the best of our knowledge, the first RhoGEF linking actin dynamics to cell cycle progression in G1. Because TEM4 activity may also be upregulated by highly dynamic F-actin (Mitin et al., 2012), and its expression is controlled by actin-regulated mechanotransduction signalling pathways such as the Hippo pathway (Lin et al., 2017; Memon et al., 2021; Y. Wang et al., 2018), we propose that TEM4 may function as a mechanosensor linking the contractile state of the cell to the cell cycle regulatory machinery in G1. In agreement with this idea, inactive or actin-binding deficient mutants of TEM4 failed to increase the mitotic index in nocodazole treated cells. Moreover, the regulation of between mechanotransduction pathways and TEM4 may be bidirectional, as suggested by loss of signalling from the Hippo pathway effector yes associated protein 1 (YAP1) after TEM4 inactivation ((Memon et al., 2021) DKP and SE, unpublished results).

TEM4 depletion resulted in a tempered cell cycle with ultimately decreased expression of the G2 and M phase cyclins and diminished CDK1-mediated phosphorylation events in the small fraction of cells that do progress towards mitosis. Consequently, TEM4-depleted cells exhibited significant defects in the execution of mitotic functions that depend on fine-tuned CDK1 activity including mitotic entry and progression, retraction fibre formation, spindle formation and positioning and accurate chromosome segregation. Although we did not find any strong defects in the SAC in cells depleted of TEM4 under the conditions tested here, the lack of sufficient cyclin B-CDK1 activity may weaken the SAC response when challenged with longer arrest times or when the SAC is further sensitized. Given the importance of cyclin B-CDK1 activity in early mitosis, it is likely that a host of additional early mitotic events are deregulated in response to TEM4 depletion.

Our observations localize the bulk of TEM4 with the cortex in mitosis although we cannot exclude the presence of a small pool at the kinetochore as previously reported (Isokane et al., 2016). Whether TEM4 contributes beyond regulation of cyclin expression to mitotic cortical functions including dissolution of focal adhesions, formation of retraction fibres and spindle positioning has not been formally ruled out. Nevertheless, we favour the hypothesis that the mitotic phenotypes observed in TEM4 depleted cells are mostly an indirect effect of the G1 delay. In agreement with this, we show that several of the mitotic defects caused by TEM4 depletion can be largely rescued by expression of exogenous cyclin B. Ultimately, cyclin B expression significantly restored mitotic progression and reduced chromosome missegregation in anaphase to control levels in agreement with the idea that most mitotic defects associated with TEM4 depletion are a result of reduced cyclin B-CDK activity. Cyclin B expression also partly rescued retraction fibres, remnants of focal adhesions that disassemble in CDK1-dependent manner before entry into mitosis (Jones et al., 2019, 2018). The rescue effect is only partial, potentially due to transgene expression at sub-endogenous levels from a single site of integration in the T-REx system. Cyclin B-CDK1 phosphorylation of MISP, a protein involved in centrosome clustering spindle orientation and positioning (Maier et al., 2013), the integrin activator kindlin (Chen et al., 2022) and the formin protein DIAPH1 (Nishimura et al., 2019) have all been shown to regulate retraction fibres and cortical tension at mitotic entry, and a recent proteomics screen has indicated that many more CDK1 targets may be able to influence this process (Robertson et al., 2015). Finally, we routinely observed that the short TEM4^1-610^ fragment localized more prominently at the cortex than TEM4^WT^ (Fig. 1B, Fig. S1A), in agreement with the idea that cortical TEM4 in mitosis may be autoinhibited through intramolecular interactions as has recent been proposed (García-Jiménez et al., 2022).

One curious observation in mitotic cells depleted of TEM4 was the hyperactivation of Rho A despite the expected decrease in contractility seen in interphase. We suspect that this might also be a result of inappropriate activation of alternative GEFs in mitotic cells as a result of lower cyclin B-CDK1 activity. In murine cells, GEF-H1 has been implicated in modulating the spindle through RhoA-DIAPH1 pathway (Bakal et al., 2005). Aurora A/B and CDK1-cyclin B phosphorylate GEF-H1 in early mitosis, thereby inhibiting catalytic activity. The RhoGEF LARG (also known as ARHGEF12) has been implicated in cytokinesis and abscission. Interestingly, LARG is also phosphorylated by CDK1 and phosphodeficient LARG is a more active GEF in cells than the phosphomimetic (Helms et al., 2016). Thus, deficient CDK1 activity may contribute to premature Rho A activation through inadequate phosphorylation of Rho GEFs normally inhibited in early mitosis. Alternatively, Rho A may be hyperactivated in mitosis as a result of decreased Rho C activity. Indeed Mitin et al. demonstrated excessive actomyosin contractility in endothelial cells depleted of TEM4 as a result of Rho A hyperactivation and proposed that TEM4 and Rho C suppress myosin contractility (Mitin et al., 2013). Further studies will be needed to define the precise contributions of Rho regulators and the different Rho isoforms at the cortex in early mitosis. Overall, our data support a model where F-actin contractility in interphase is regulated by TEM4 to drive timely progression of the cell cycle at the G1/S transition.

## Materials and Methods

### Cell lines

HeLa-T-REx Flp-IN and HCT-116 cell lines were cultured at 37°c with 5% CO2 in Dulbecco’s Modified Eagle Medium (DMEM, Hyclone) and RPMI medium, respectively, supplemented with 10% fetal bovine serum and 1% Penicillin-Streptomycin (100 µg/ml-1, Hyclone). For lentiviral induction, TEM4-21 and TEM4-22 cells were treated with 0.5 µg/ml of Doxycycline for 48 or 72 h as specified.

### Drug treatments and transfections

Drug treatment was performed as follows, unless otherwise indicated: Thymidine (Acros Organics, 2mM for 16 h), MG132 (Calbiochem, 20μM for 2.5 h), nocodazole (Sigma,100ng/mL for 12 h), S-trityl-L-Cysteine (STLC, Sigma, 5 μM for 12 h), Paclitaxel (taxol, Calbiochem, 15nM for 14 h), Y27632 (Calbiochem 10 μM for 30 mins). Plasmid transfections were performed using FuGENE® HD according to manufacturer’s protocol.

### ShRNA construction and generation of inducible cells lines and shRNA TEM4 resistant cell lines

HeLa-TREx stable cell lines expressing inducible shRNA targeting TEM4 (TEM4-21 and TEM4-22) have been previously described (Prifti et al., 2022). The same approach was used for generation of the HCT116 lines with inducible TEM4 depletion. After lentivirus preparation and infection, single cell clones were expanded and screened via immunoblot and genomic sequencing. HeLa-TREx stable cell lines expressing TEM4^WT^, TEM4^ΔABD^ and TEM4^Y1216A^ were generated using Flp-In system (Life Technologies) according to the manufacturer’s instructions. TEM4 cDNA was cloned in pcDNA5-FRT-TO with and N-terminal 3xMYC-GFP tag and was rendered resistant to both sh21 and sh22 by introducing silent mutations using the following primers: shRNA-TEM4-21 Forward 5’- GTACCTCAATAATCAAGTATTTGTGTCTCTGGCCAATGGAGAG-3’, Reverse 5’ ATACTTGATTATTGAGGTACAAGATGCAGGTCACAGAGGCCG-3’ and for shRNA-TEM4-22 Forward 5’- AATACTCTGGATCACCTCCTGATATCCCTTGCTCTCTTCACTGC-3’ and Reverse 5’-AGGTGATCCAGAGTATTGTTCAGGGGCCTGGCACCCTGGGGCGTG-3’. As both shRNA and rescue constructs are under control of the doxycycline promoters, expression of both is achieved simultaneously.

### Microtubule regrowth assay

Microtubule regrowth was performed as previously published (Tulu et al., 2006). Essentially, cells were plated and two days later they were treated with 3.3 μM of nocodazole for 3h. Media containing nocodazole was washed out 4 times with warm PBS, followed by a wash in warm media. Cells were then incubated in warm media as indicated to allow the microtubules to regrow. Cells were fixed with cold methanol for 10 minutes at – 20 °C for simultaneous fixation and permeabilization.

### Cold treatment assay

Cells were washed twice with cold PBS and incubated on ice for 20 mins in the presence of cold DMEM medium. Fixation was performed with cold PTEMF buffer (0.2% Triton X-100, 20mM PIPES pH 6.8, 1mM MgCl2, 10mM EGTA and 4% formaldehyde) for 10 mins.

### Protein extraction and Western blotting

Cell pellets were lysed in RIPA lysis buffer (150mM Tris-HCL pH 7.5, 150mM NaCl, 10mM NaF, 1% NP-40, 0.1% Na-deoxycholate) or 8M urea lysis buffer (8M urea, 50 mM HEPES pH 7,5, 5% glycerol, 1.5mM MgCl_2_ and the same protease and phosphatase inhibitor cocktail used in the RIPA buffer) and a protease and phosphatase inhibitor cocktail that included 20mM B-glycerophosphate, 0.1mM sodium vanadate, 10mM sodium pyrophosphate, 1mg.ml-1, leupeptin, 1mg.ml-1, aprotinin and 1mM AEBSF. Lysed cells were incubated on an orbital shaker at 4°C for minimum 30 minutes and followed by centrifugation at maximum speed for 20 mins at 4°C. The supernatant was collected and quantified using the BCA assay (Thermo Scientific) prior to Western blotting. Protein extracts were diluted in SDS-PAGE sample buffer and loaded onto 8-12% SDS-PAGE gels. Membranes were blocked in 5% of milk diluted in PBS and containing 0.05% of Tween-20 for 30 minutes, and then, incubated overnight at 4°C with primary antibody in 5% of milk in PBS. Membranes were washed with PBS containing 0.05% of Tween-20 and incubated with the appropriate secondary antibody conjugated to horseradish peroxidase for 1 h at room temperature. After three additional washes, antibody binding was detected with either the Clarity or Clarity Max Western ECL substrate (Bio-Rad) and ChemiDoc MP Imaging System (Bio-Rad). The following antibodies were used for Western blotting: The anti-TEM4 used is a custom-made Rabbit polyclonal antibody (Biomatik) generated against the following peptide sequence: RWSGGPGLREEDTDTPG-Cys. Peptide purified serum was subsequently used for Westerns; anti-Cyclin A (clone 25, BD Transduction Laboratories); anti-Cyclin B1 (V152, Cell signaling); anti-Cyclin D1 (92G2, Cell Signaling); anti-Cyclin E (HE12, sc-247, Santa Cruz); anti-GFP (1:500, Millipore); Anti-pSer/Thr-Pro MPM-2 (EMD Millipore), anti-γ-tubulin (TU-30, ab27074, Abcam); anti-GAPDH (1A10, NBP1-47339, Novus Biologicals); and anti-NuMA (GT3611, Thermo Fisher Scientific).

### RNA extraction and purification

Cell pellets were incubated with RLT lysis buffer (Qiagen) containing 10% β-mercaptoethanol (Sigma, M3148) for total RNA extraction which was further purified with the RNeasy Mini Kit (Qiagen) according to the manufacturer’s protocol. Samples were incubated with RNase-free DNase (Qiagen) to eliminate potential genomic contamination and Nanodrop 1000 microvolume spectrophotometer (Thermo Fisher Scientific) was used to evaluate the quantity and quality of purified total RNA. Samples were stored at −80°C until use.

### Reverse transcription and quantitative real-time PCR (qRT-PCR)

iScript Advanced cDNA Synthesis Kit for PCR (BioRad) was used for Reverse Transcription (RT) of 1µg for each sample, following the manufacturer’s protocol. Real-time quantitative PCR was carried out on cDNA samples using SsoAdvanced Universal SYBR Green Supermix (BioRad). The primer sequences were as follows: *HPRT1*: forward 5’-TGCTGAGGATTTGGAAAGGGT-3’, reverse 5’- AGCAAGACGTTCAGTCCTGT-3’; *ARHGEF17*: forward 5’- TACATGCTGAACCTGCACTCC-3’, reverse 5’-GTGCTTCCGCATGTCCACC-3’; *CCNB1*: Forward 5’-GATACTGCCTCTCCAAGCCC-3’, reverse 5’- TTTCCAGTGACTTCCCGACC-3’; *CDKN1A*: forward 5’- AGACTCTCAGGGTCGAAAACG-3’, reverse; 5’-ATGCCCAGCACTCTTAGGAA-3’; *CCNE1*: forward 5’-AGAGGAAGGCAAACGTGACC-3’, reverse 5’- GAGCCTCTGGATGGTGCAAT-3’; 0.5 μM of forward and reversed primer were combined with 2 μL cDNA samples, for the qRT-PCR reaction. Two negative controls were included: an RT-negative control and a no-template control. Samples were incubated at 95°C for 5 minutes followed by 40 cycles of three amplification steps: 95°C for 15 seconds, the optimal primer-specific temperature (between 50 and 60°C) for 15 seconds and 72°C for 15 seconds. Each qRT-PCR reaction was performed as two technical replicates for each biological sample, and then, normalized to HPRT. The Pfaffl method was used to analyze the results (Pfaffl, 2001). Fold inductions were calculated, and the results were expressed as relative quantification values based on cycle threshold (Ct) comparisons between different samples. Experiments were performed on 3-6 biological replicates as indicated.

### Immunofluorescence

Cells were grown on coverslips coated with 25µg/ml poly-ethylenimine (PEI) in 150 mM NaCl and fixed in PTEMF buffer (0.2% Triton X-100, 20mM PIPES pH 6.9, 1mM MgCl_2_, 10mM EGTA and 4% formaldehyde) for 10 minutes at room temperature. For immunofluorescence of endogenous MPS1, cells were incubated for 5min with 1% formaldehyde in PBS, quenched in 0.1 M glycine for 1h, then permeabilized in 0.1% Triton X-100 for 3min. For pERM visualization cells were fixed with 10% trichloroacetic acid (#T0699; Sigma-Aldrich) for 10 mins at room temperature before extensive washings with TBS as previously described (Carreno et al., 2008). After fixation, coverslips were blocked with 3% bovine serum albumin (BSA) in PBS-Tween 0.2% for at least 30 minutes prior to incubation with primary and secondary antibodies for 2 h and 1 h, respectively, at room temperature. Antibodies were used at 1 µg/ml, unless otherwise indicated. Antibodies against the following proteins were used: α-tubulin (DM1A, Santa Cruz); CREST HCT-0100, Immunovision); MPS1 (M5818, Sigma-Aldrich); Rho A-GTP (NewEast biosciences); GFP (mix of clone 7.1 and 13.1, Roche) and α-tubulin Alexa-647 (11H10,Cell Signaling); Myc (9E10, Fisher) and H3pT3 (SB153a, Southern Biotechnology); Cyclin A (clone 25, BD Transduction Laboratories) Cyclin B1 (D5C10, Cell Signaling); BUBR1 (Elowe et al., 2007); pERM (gift of S.Carréno). HOESCHT 33342 (Sigma) was used to stain chromatin and phalloidin–Atto 565 (9402, Sigma Aldrich) for F-actin. Alexa Fluor-affiniPure series secondary antibodies (Thermo) or (Jackson ImmunoResearch) were used for immunofluorescence (1:1000). For live-cell imaging, cells were washed with PBS and treated with SiR-DNA dye (CY-SC007, Cytoskeleton) and SPY555-Tubulin (CPY-SC203, Cytoskeleton) probe at 300 nM for 1h prior imaging.

### Flow Cytometry Analysis

Cells were treated with 0.5 µg/ml of doxycycline for 72 h and synchronized with 3.3 uM nocodazole for 12 h. Cells were then washed with PBS and 2×10^6^ cells/mL were incubated at 37°C for 60 mins with complete medium containing 5 μg/mL Hoechst 33342. Cells were then pelleted by centrifugation and the medium containing Hoechst 33342 was aspirated. Cells were washed in 1X PBS and flow cytometry analysis followed with a BD LSRII SORP flow cytometer (Becton Dickinson, New Jersey, USA) equipped with BDFACSDiva software. Hoechst dye was excited with a 355 nm UV laser emitted fluorescence and detected with a 450/50 bandpass filter. Cell cycle analysis was performed using FlowJo software v10 (Becton Dickinson) and the Watson pragmatic algorithm (Watson et al., 1987).

### Confocal Microscopy

An inverted Olympus IX80 microscope equipped with a WaveFX-Borealin-SC Yokagawa spinning disc (Quorum Technologies) and an Orca Flash4.0 camera (Hamamatsu) was used for the acquisition of all images. Images were acquired by Metamorph software (Molecular Devices). Optical sections with identical exposure times for each channel within an experiment were acquired and then projected into a single picture using ImageJ (Fiji version 1.52i). The system was equipped with a motorized stage (ASI) and incubator with atmospheric CO_2_ warmed at 37°C for live-cell imaging. For time-lapse experiments, images were taken with a 20X objective every 3 minutes.

### Quantitative polarization microscopy

Cell contractility was measured with quantitative polarization microscopy (qPOL) as described previously (W. Wang et al., 2018). Images were acquired using a 40× polarization lens (NA=0.9) with 10° intervals of the polarizer rotation over a range of 0 to 180°. The polarized image sequences were processed with a custom MATLAB code to obtain an optical retardance map. The retardance signal proportional to cell contractility was quantified by measuring the average retardance over the cell area after background subtraction in ImageJ2.

### Image quantification and statistical analysis

Unless otherwise stated, all experiments and statistical analyses were performed in biological triplicates. Image J2 was used for image processing and all images shown in the same figure have been identically scaled. For measurement of signal intensities at the kinetochores, cells were measured individually by cropping the desired cell, creating a selection of the kinetochores area by thresholding, and transferring to and measuring the selection in the desired channels and intensities were measured according to established protocol (Saurin and Kops, 2016) in ImageJ2 (Fiji version 1.52 i) (Rueden et al., 2017; Schindelin et al., 2012). For whole spindles, intensity measurements with or without cold treatments were performed according to a published protocol (DeLuca et al., 2016). Integrated signal intensity was measured in all relevant channels and intensities indicated are values relative to a tubulin, CREST or arbitrary units. To determine the spindle angles (α), z-stack images of metaphasic cells immunostained with anti γ-tubulin (spindle poles) and anti α-tubulin antibodies (mitotic spindle) were acquired. Spindle angles were calculated in Image J as previously published (Cau et al., 2013; Mannen et al., 2016). Once angles were calculated, negative angles or angles above 90° were transformed to the first quadrant. Angular histograms were plotted using a custom R script (Scarpa et al., 2018). Misaligned and lagging chromosomes were quantified in fixed cells as previously described (Gama Braga et al., 2021). Briefly, spindle pole position was defined, followed by the establishment of an alignment region after equally dividing the area in between the poles of the cell into 5 zones. The alignment zone was defined as the middle zone and was excluded from further analysis. The number of misaligned kinetochores lying outside were considered misaligned at metaphase and were counted.

Identification of lagging chromosomes in anaphase cells followed the same principle as above where the coordinates at the centroids of the two anaphase chromosome masses were selected and the region of interest in between was defined. Kinetochores inside this region were considered lagging and were quantified. Cortical intensity was measured in a single confocal z medial slice in Image J2 using a freely drawn 5 pixels line. Mean pixel intensity was measured, and individual values were normalized to control cells. Cell rounding was measured in Image J using the “fit ellipse” tool. The circularity index *C* (= 4π(area/perimeter^2^)) was used to quantify the roundness of a two-dimensional object corresponding to the cross-section of the cell by a plane.

### Statistical analysis

Statistical analysis and graphic plotting were performed in GraphPad Prism Software V6.01 and presented as Superplots where applicable (Lord et al., 2020). A non-parametric t-test (Mann-Whitney test) was used in experiments comparing only two conditions, and ANOVA (Kruskal-Wallis test) one-way or two-way was used for experiments with more than two conditions. Statistical significance was considered with a P-value smaller than 0.05 and all P-values are reported.

## Supporting information

Control Mitosis

Mitosis TEM4_SH22

Mitosis TEM4_SH21

## Acknowledgements

The authors thank Sébastien Carréno for the pERM antibodies, Elena Scarpa for the gift the spindle angle normalisation macro, Christian Doucet and Vincent Desrosiers for technical assistance, and Jan Ellenberg for helpful discussions. Work in the Elowe lab is supported is supported by Canadian Institutes of Health Research grants to S.E. (156405, 153046) and the national science and engineering council of Canada (RGPIN-2016-05841). Work in the Bordeleau lab is supported by the national science and engineering council of Canada Discovery grant to F.B. (RGPIN-2018-06214). S.E. holds an FRQS (Fonds de Recherche de Santé Québec) Senior researcher salary award. D.K.P has been supported by training awards from Desjardins and by a CRCHU de Québec training award. F.B. is a tier 2 Canada Research Chair in Tumor Mechanobiology and Cellular Mechanoregulation.

## SUPPLEMENTAL FIGURE LEGENDS

**Figure S1:**
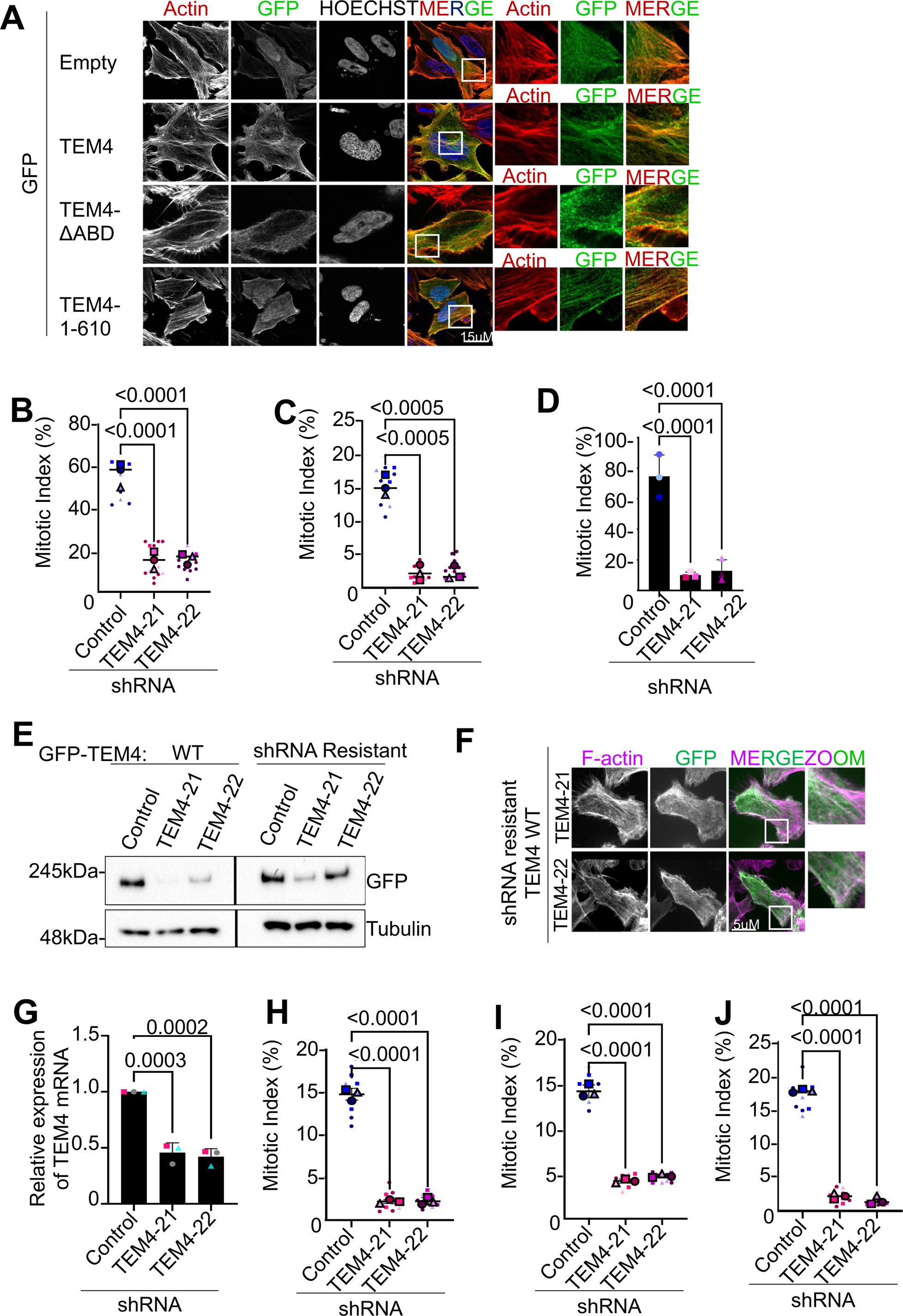
TEM4 decreases mitotic index in different cell lines independently of the microtubule’s status. A: HeLa-T-REx were transfected with indicated TEM4 constructs before being fixed. Cells were stained with anti-GFP (green), Phalloidin-Atto 565 for F-actin (red) and Hoechst (blue). Scale bar = 15μm. B: Control or HeLa-TREx cells depleted of TEM4 were treated with 15nm taxol for 12 h and fixed. Data shown is mean ± SEM (from ∼1400 cells, n=3). C: Control or HeLa-TREx cells depleted for TEM4 were synchronized in mitosis after 10 h of release from double thymidine block. Data shown is mean ± SEM (from ∼1200 cells, n=3). D: Control or HeLa-TREx cells depleted for TEM4 were released from double thymidine and treated with MG132 at mitotic entry. Data shown is mean ± SEM (from ∼1500 cells n=3). E: Expression of shTEM4-21 and shTEM-22 resistant TEM4 versus TEM^WT^ proteins in control, TEM4-21 and TEM-22 cell lines. Lysates were blotted for the indicated antibodies. F: Expression and localization of shRNA resistant TEM4^WT^ versus TEM^WT^ proteins in TEM4-21 and TEM-22 cell lines. Cells we stained with anti-GFP and Phalloidin-Atto 565. Scale bar = 5μm. G: Expression of *ARHGEF17* mRNA in HCT-116 cells transduced with shTEM4-21 or shTEM4-22. ShRNA expression was induced for 72 h using 0.5 μg/ml doxycycline. H: Mitotic index in HCT116 cells expressing shTEM4-21 or shTEM4-22. ShRNAs were induced for 72 h and released for 10 h from double a thymidine block. Data shown is mean ± SEM (from ∼1200 cells, n=3). I: Mitotic index in HCT116 cells expressing shTEM4-21 or shTEM4-22. ShRNAs were induced for a total of 72 h and cells were treated with 15nm taxol for 12 h and fixed. Data shown is mean ± SEM (from ∼1000 cells, n=3). J: Mitotic index in HCT116 cells expressing shTEM4-21 or shTEM4-22. ShRNAs were induced for a total of 72h and cells were released from double thymidine block and treated with the proteosome inhibitor MG132 to prevent cells from exiting mitosis. Data shown is mean ± SEM (from ∼1500 cells, n=3).

**Figure S2:**
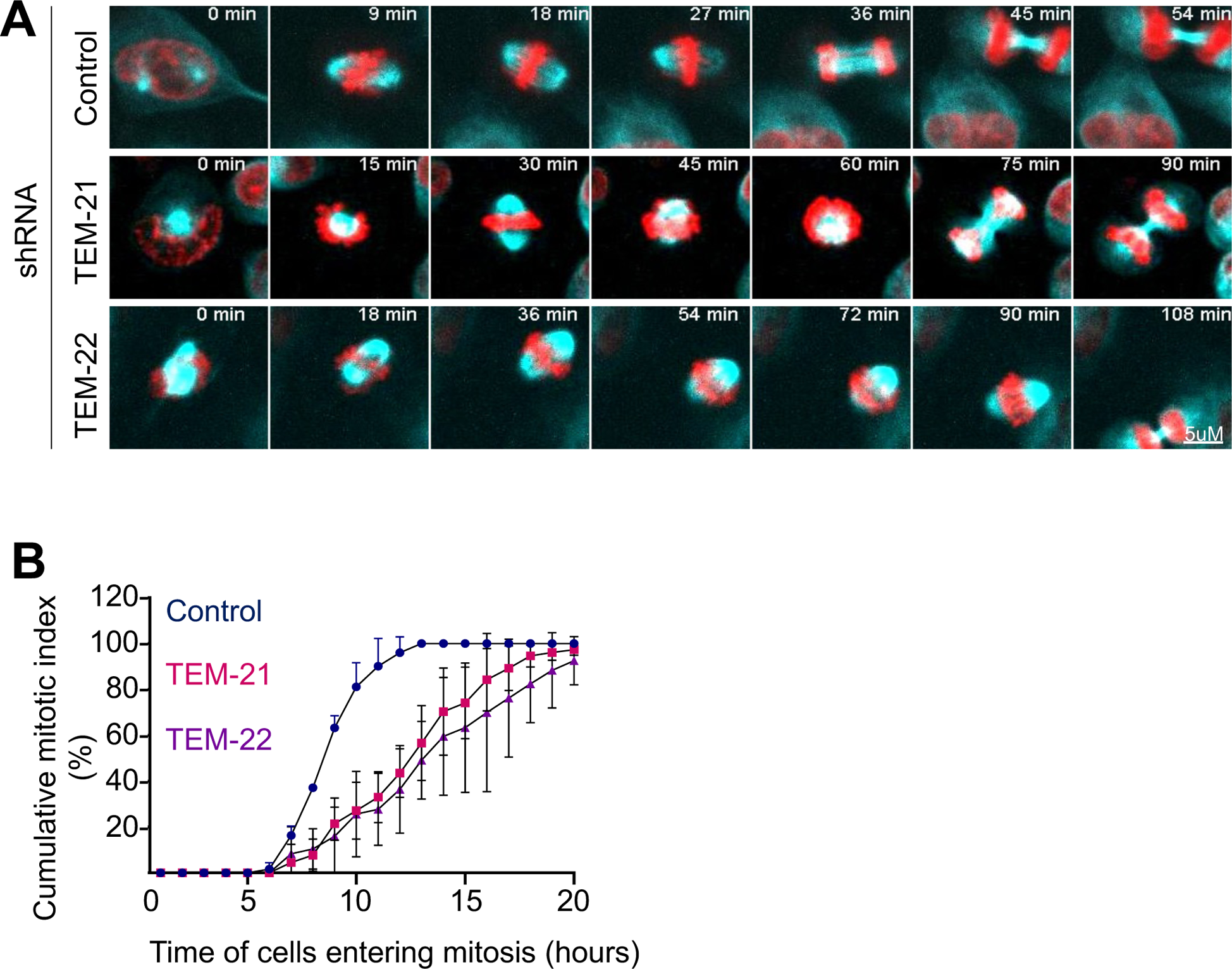
Delayed mitotic entry and in the absence of TEM4. A: Representative stills of time-lapse movies of control, shTEM4-21 and shTEM4-22 cells 72 h after induction. For imaging, cells were synchronized in mitosis after release from double thymidine block and incubated with SiR-DNA and SPY555-Tubulin to visualize the chromatin and the mitotic spindle. B: The cumulative mitotic index was measured from time-lapse movies of HeLa-T-Rex cells depleted of TEM4 shown in A. Data shown is mean ± SEM (from ∼300 cells, n=3).

**Figure S3:**
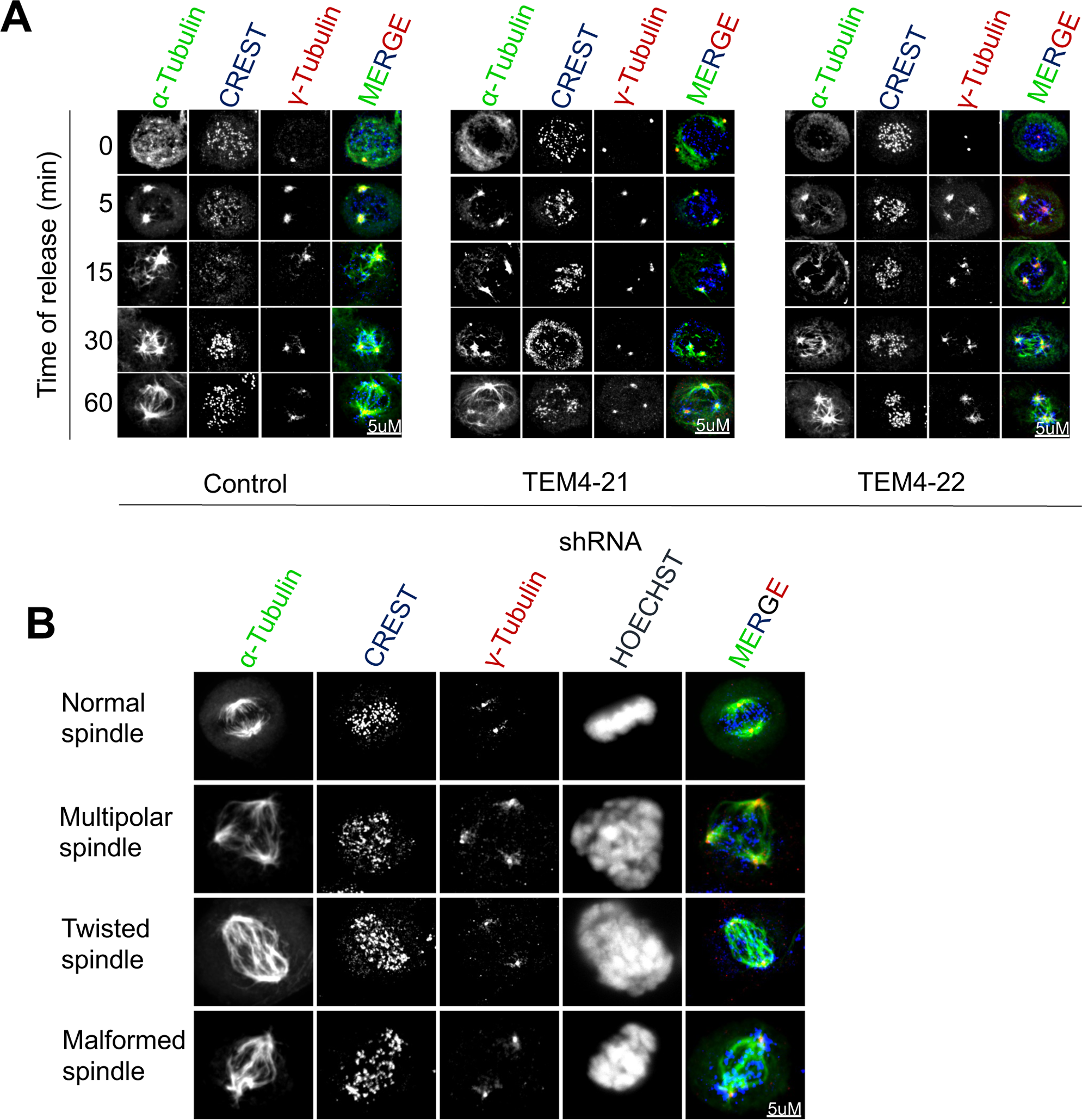
TEM4 depletion slows microtubule regrowth and results in spindle. A: HeLa-T-Rex control or TEM4-depleted cells were treated for 3 h with 3μM of nocodazole. Nocodazole was washed out and cells were incubated with fresh media for the indicated time points. Cells were fixed with PTEMF buffer and stained with CREST (blue), anti-α-tubulin (green) and anti-γ-tubulin (red). Scale bar = 5μm. B: Cells treated as in A. Representative images of mitotic spindle phenotypes observed after spindle reformation.

**Figure S4:**
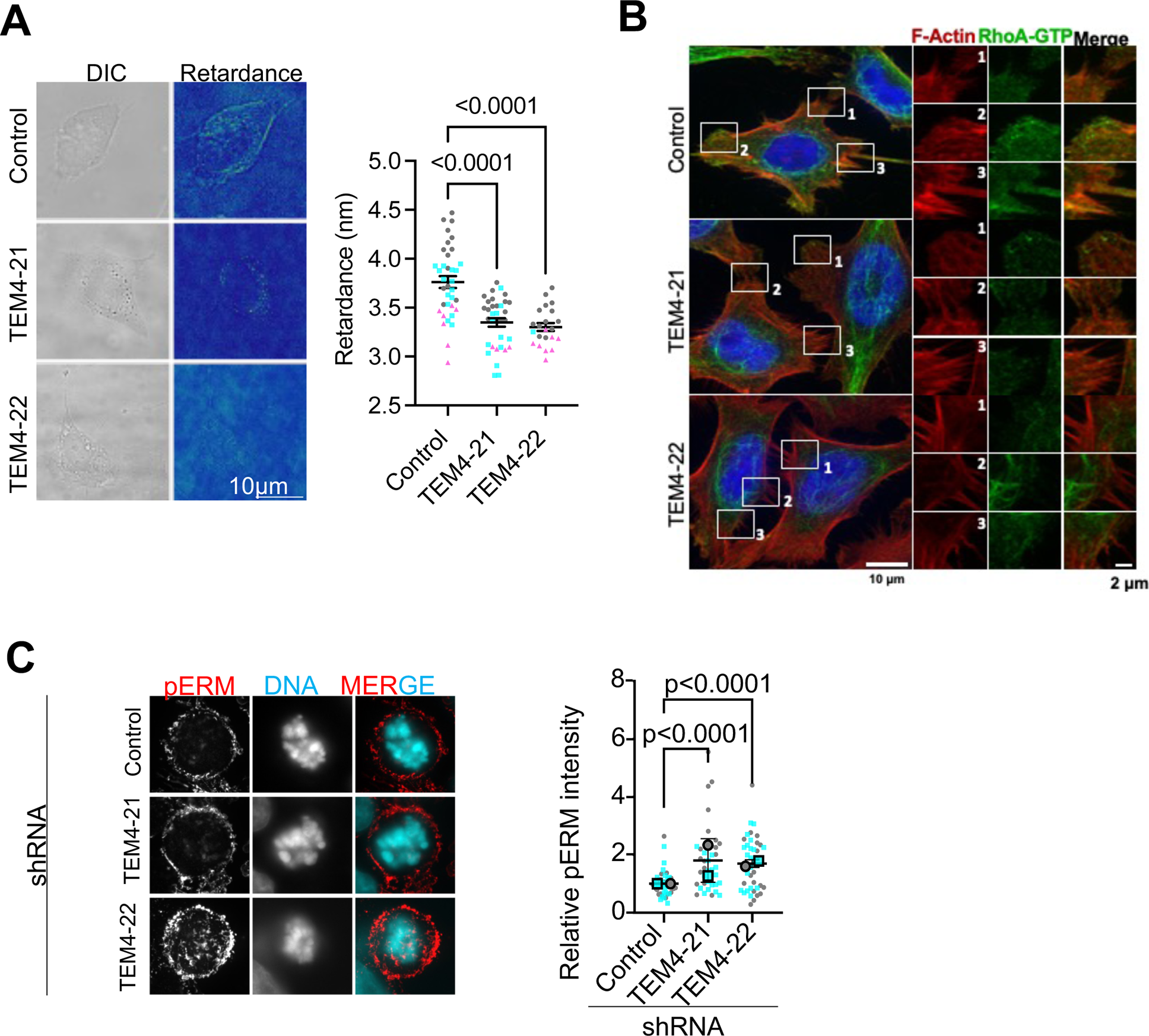
measurement of contractility markers in control and TEM4-depleted cells. A: Polarization microscopy of control and TEM4 depleted cells in interphase. Representative images and quantification optical retardance, n=3. B: RhoA-GTP staining in control and TEM4-depleted cells. Fixed cells were stain with Phalloidin-Atto 565 (red), RhoA-GTP (green) and Hoechst (blue). C: pERM staining in control and TEM4 depleted cell. Representative IF images of cells stained with pERM (red) and Hoescht (blue) Data shown is mean ± SD, n=2.

**Figure S5:**
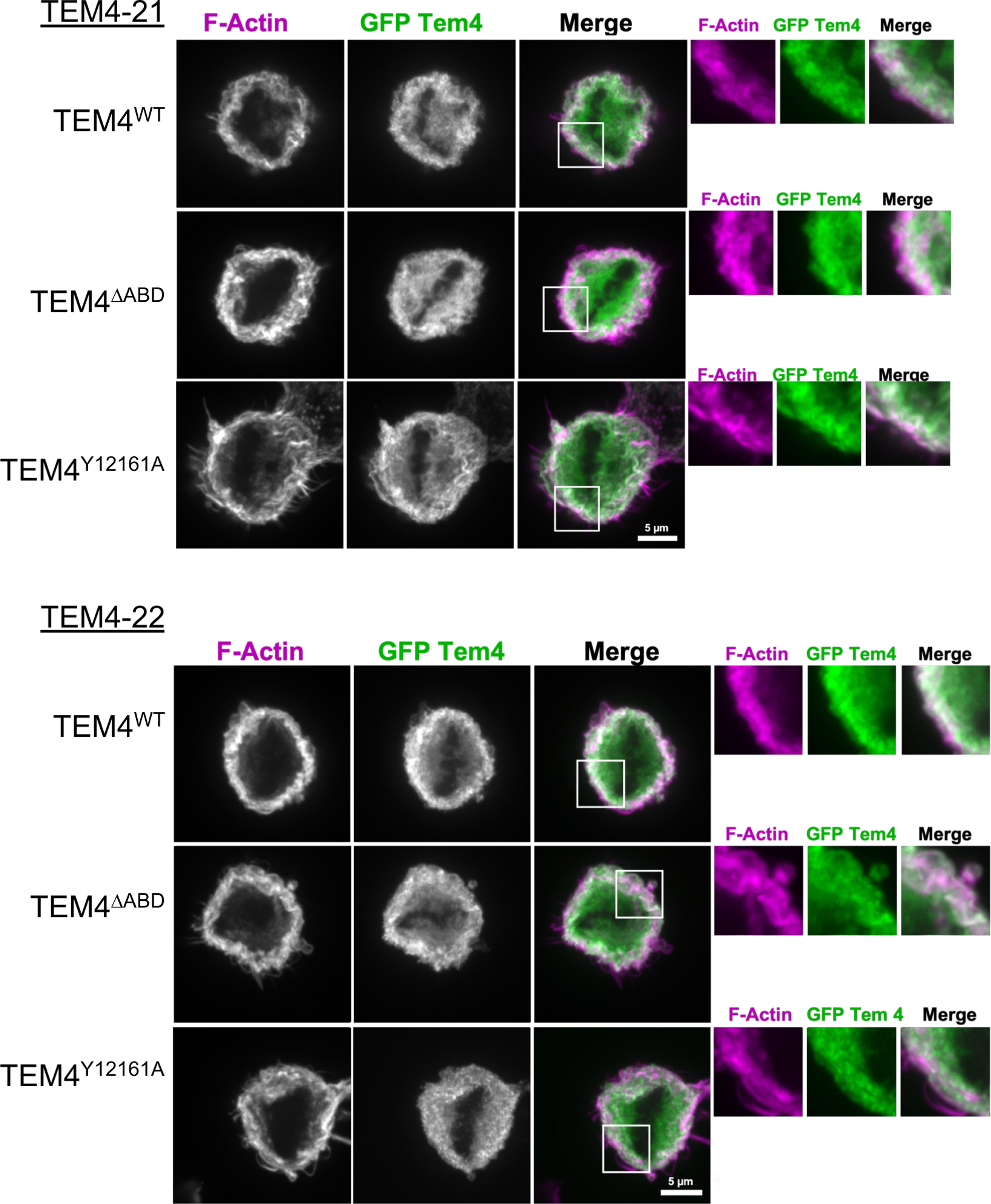
Generation of stable cell lines expressing shRNA TEM4 constructs. Immunfluorescence images of stable cell lines expressing shResistant TEM4^WT^, TEM4^1′ABD^ and TEM4^Y1216A^.

**Figure S6:**
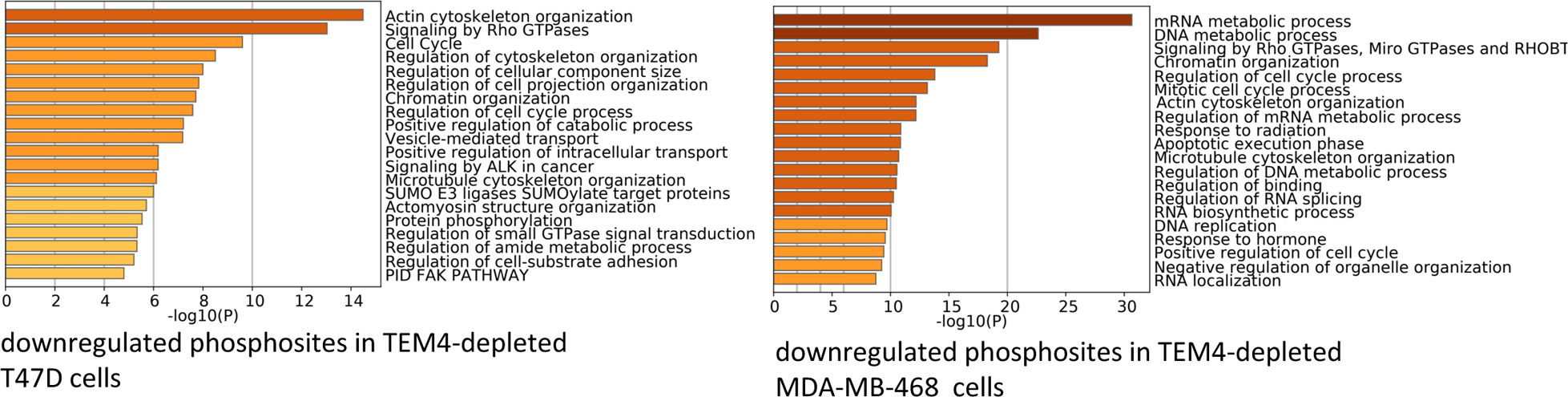
Enchrichment of GO annotation terms in phosphosites identified after depletion of TEM4. Gene ontology analysis performed with Metascape on the list of hypophosphorylated protein in T47D and MDA-MB-468 cells depleted of TEM4 as identified by Memon *et al*. The full list of genes is provided in supplementary data 1.

**Supplemental table 1**: Hyper- and hypo-phosphorylated sites in TEM4-depleted T47D and MDA-MB-468 cells.

**Supplemental table 2**: Enrichment of Ser-Pro and Thr-Pro motifs in TEM4 amongst hypo-phosphorylated sited in TEM5-depleted T47D and MDA-MB-468 cells.

**Supplemental movie 1**: Example mitosis from control cells.

**Supplemental movie 2**: Example mitosis from cells depleted of TEM4 with TEM4-21.

**Supplemental movie 3**: Example mitosis from cells depleted of TEM4 with TEM4-22.

## Notes

### Competing Interest Statement

The authors have declared no competing interest.

